# Dissociation of β_2_m from MHC Class I Triggers Formation of Noncovalent, Transient Heavy Chain Dimers

**DOI:** 10.1101/2021.07.02.450866

**Authors:** Cindy Dirscherl, Sara Löchte, Zeynep Hein, Janine-Denise Kopicki, Antonia Regina Harders, Noemi Linden, Andreas Karner, Johannes Preiner, Julian Weghuber, Maria Garcia-Alai, Charlotte Uetrecht, Martin Zacharias, Jacob Piehler, Peter Lanzerstorfer, Sebastian Springer

## Abstract

At the plasma membrane of mammalian cells, major histocompatibility complex class I molecules (MHC-I) present antigenic peptides to cytotoxic T cells. Following the loss of the peptide and the light chain beta-2 microglobulin (β_2_m), the resulting free heavy chains (FHCs) can associate into homotypic complexes in the plasma membrane. Here, we investigate the stoichiometry and dynamics of MHC-I FHCs assemblies by combining a micropattern assay with fluorescence recovery after photobleaching (FRAP) and with single molecule co-tracking. We identify non-covalent MHC-I FHC dimers mediated by the α_3_ domain as the prevalent species at the plasma membrane, leading a moderate decrease in the diffusion coefficient. MHC-I FHC dimers show increased tendency to cluster into higher order oligomers as concluded from an increased immobile fraction with higher single molecule co-localization. In *vitro* studies with isolated proteins in conjunction with molecular docking and dynamics simulations suggest that in the complexes, the α_3_ domain of one FHC binds to another FHC in a manner similar to the β_2_m light chain.

**Significance Statement:** MHC class I molecules are cell surface transmembrane proteins with key functions in adaptive immunity against viral infections. The spatiotemporal organization of fully assembled MHC I at the cell surface and its function with respect to trans-interactions with T and NK cells has been studied in detail. By contrast, the consequences of peptide and β_2_m dissociation yielding to formation of free heavy chains (FHC) have remained unclear. We have discovered that class I free heavy chains form distinct non-covalent dimers at the cell surface rather than non-specific clustering, and we have identified a dimerization interface mediated by the α_3_ domain. We propose that these non-covalent dimers are the basis of distinct signaling and endocytic sorting of MHC I FHC. This is to be explored in further work.

## Introduction

Major histocompatibility class I molecules (MHC-I) fulfil central tasks of the adaptive immune response against infections and malignancies by presenting antigenic peptides to T cells (Comber and Philip, 2014; Kaufman, 2018; Townsend and Bodmer, 1989). MHC-I heterotrimers consist of the polymorphic transmembrane heavy chain (HC), the non-polymorphic soluble light chain beta-2 microglobulin (β_2_m), and the peptide (Townsend et al., 1990, 1989). In addition to this trimer, two more states of MHC-I occur at the cell surface, the ‘empty’ HC/β_2_m heterodimer that lacks peptide (Ljunggren et al., 1990; Sebastián Montealegre et al., 2015), and the monomeric ‘free’ heavy chain (FHC) (Edidin et al., 1997; Geng et al., 2018). Since the binding of peptide and β_2_m is cooperative (Elliott et al., 1991; Gakamsky et al., 1996), HC/β_2_m heterodimers are conformationally unstable, and loss of peptide leads to the rapid formation of FHCs (schematic in **Figure 1A**) and to the subsequent endocytic removal of FHCs by a sorting mechanism that is not understood (Sebastián Montealegre et al., 2015). For FHCs present at the cell surface, important regulatory functions mediated by homo- and heteromeric interactions in *cis* and *trans* have been proposed (Arosa et al., 2021, 2007; Campbell et al., 2012), which suggest defined spatiotemporal organization and dynamics of FHC in the plasma membrane. Indeed, clustering and covalent dimerization of MHC-I have been identified using a variety of approaches including recombinant proteins and live cells (Allen et al., 1999; Antoniou et al., 2011; Armony et al., 2021; Baia et al., 2016; Blumenthal et al., 2016; Bodnar et al., 2003; Capps et al., 1993; Chakrabarti et al., 1992; Fassett et al., 2001; Ferez et al., 2014; Lu et al., 2012; Makhadiyeva et al., 2012; Matko et al., 1994; Triantafilou et al., 2000); these represent complexes of different and largely unclear composition, size, and type of intermolecular bonding.

**Figure 1:**
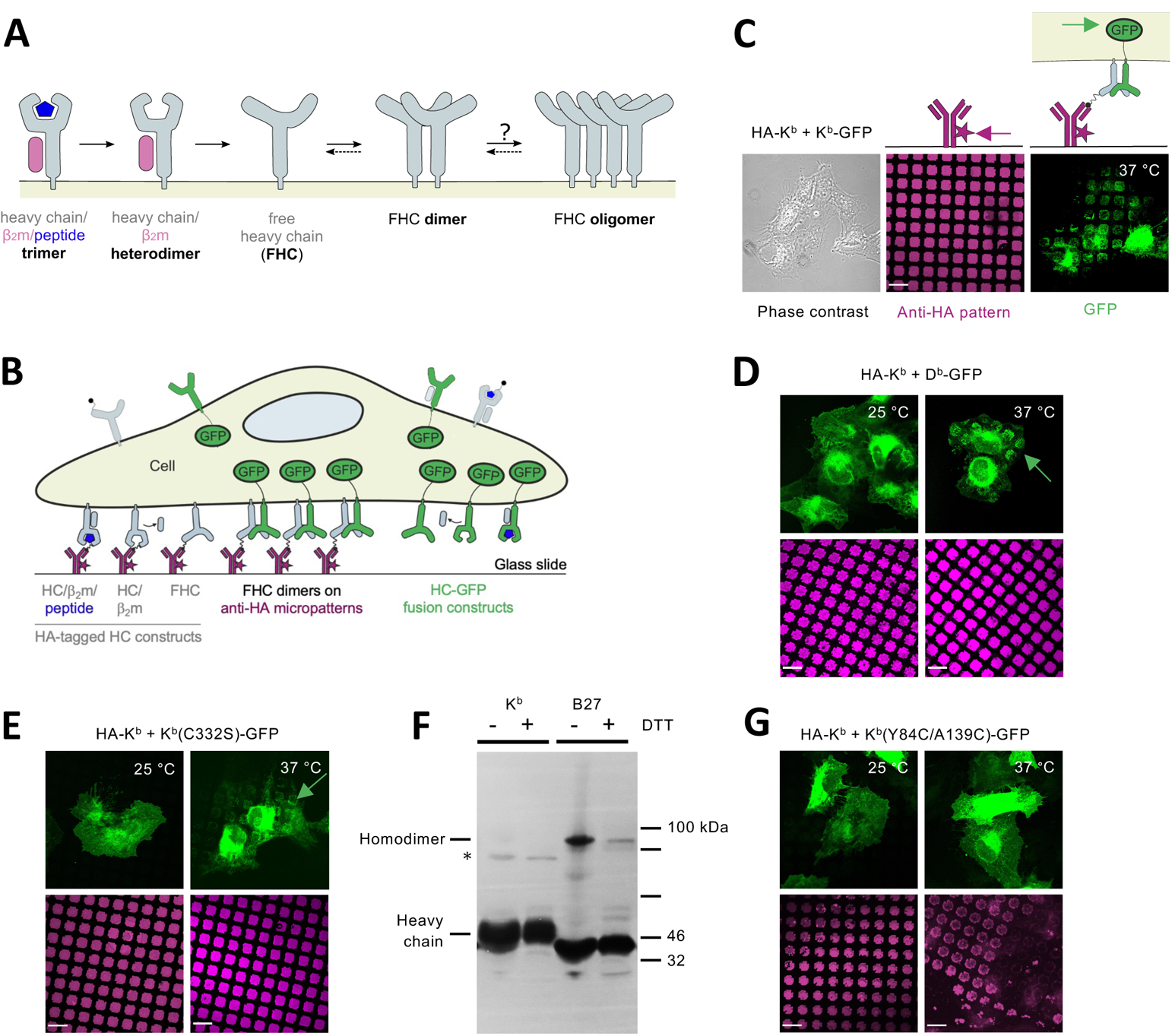
MHC I HC/HC association requires dissociation of β_2_m but no disulfide bonds. **A**, Schematic of MHC class I states at the plasma membrane. Dissociation of peptide from the HC/β_2_m/peptide trimer results in an ‘empty’ HC/β_2_m heterodimer. Dissociation of β_2_m then produces free HCs (FHCs), which can form FHC associations (HC/HC dimer and oligomer shown). Other forms such as disulfide-linked dimers are known depending on the allotype (see the text). **B**, Schematic representation of the two-hybrid antibody micropattern assay. Cells expressing a class I GFP fusion (green) and an N-terminally HA-tagged class I (gray) are seeded onto glass slides that are printed with micrometer-sized patterns of fluorescently labelled anti-HA antibodies. Dissociation of β_2_m generates FHCs of both constructs, which diffuse freely in the plasma membrane and eventually associate with each other to form HC/HC dimers (center) or oligomers (not shown). The HC/HC associations, which contain both HA-tagged and GFP-fused FHCs, localize in the pattern elements and are visible as pattern-shaped GFP fluorescence on the plasma membrane. **C**, Representative fluorescence micrograph showing one single STF1 cell expressing both HA-tagged and GFP-fused H-2K^b^ in phase contrast (left), the purple anti-HA antibody pattern on the glass slide (middle), and green K^b^-GFP colocalizing with the antibody pattern (right). The arrows point to the fluorophore observed in the respective panel and emphasize the plane of the image. Scale bar, 20 μm. **D**, Interaction occurs between HA-tagged K^b^ and D^b^-GFP FHCs (37 °C) but not between HC/β_2_m heterodimers (25 °C). Scale bar, 20 μm. **E**, HC/HC interaction does not involve intracellular disulfide bond formation, since the FHCs (37 °C) of HA–K^b^ and K^b^(C332S) GFP, which lacks the cytosolic cysteine, interact in the micropattern assay. Scale bar, 20 μm. **F**, B27, but not K^b^, forms covalent dimers. HA-K^b^ (K^b^ in the label) and HA-B*27:05 (B27) molecules were immunoprecipitated from the lysate of transduced STF1 cells with an anti-HA monoclonal antibody, separated by reducing (+) or nonreducing (–) SDS-PAGE, and monomers (heavy chain) and covalent homodimers as indicated were detected by Western blotting with an anti-HA antiserum. * denotes a background band. **G**, No interaction of K^b^–GFP with F pocket-stabilized HA–K^b^(Y84C/A139C), a disulfide-stabilized K^b^ variant with increased β_2_m affinity. Scale bar, 20 m.

Recently, we have achieved direct detection of homomeric FHC interactions in the intact plasma membrane by means of a live-cell two-hybrid micropattern assay (**Figure 1B, C**) (Dirscherl et al., 2018): micrometer-sized patterns of anti-hemagglutinin tag (HA) monoclonal antibody are printed onto glass coverslips (Schwarzenbacher et al., 2008; Sevcsik et al., 2015). Onto these micropatterns, cells are seeded that express two different MHC-I HC constructs: one construct has an N-terminal (extracellular) HA tag and thus, while diffusing laterally in the plasma membrane, is captured into the printed antibody pattern. The other has no HA tag but a C-terminal (intracellular) green fluorescent protein (GFP) fusion domain (**Figure 1B**). Interaction of GFP-tagged HC with micropatterned HA-HC is detected by an increased GFP fluorescence of the pattern elements (**Figure 1C**). The system allows the observation of different defined conformational states of class I: with TAP2 (transporter associated with antigen processing)-deficient fibroblasts (which cannot transport peptide into the endoplasmic reticulum (ER)), empty HC/β_2_m heterodimers are present at the cell surface. By adding peptide, shifting cells to 25 °C, or incubating at 37 °C, we therefore can accumulate trimers, HC/β_2_m heterodimers, or FHCs, respectively, at the cell surface (**Figure 1B** and described below). These experiments revealed that formation of homomeric MHC-I only takes place in the absence of β_2_m, *i.e.*, only between FHCs.

We now have uncovered the molecular principles that govern such homomeric MHC-I FHC association in the plasma membrane. Cell micropatterning in conjunction with fluorescence recovery after photobleaching (FRAP) was used to probe dynamics, stability, and prominence of FHC complexes. We directly demonstrate FHC association in the plasma membrane under physiological conditions by using real-time single-molecule tracking (SMT) and co-tracking (SMCT). Surprisingly, we find that FHCs transiently associate into non-covalent dimers with lifetimes in the sub-second range. Based on our findings that the HC/HC complexes contain no β_2_m and that the α_3_ domain of the HC is sufficient for dimerization *in vitro* and in cells, we propose a molecular model structure of MHC-I HC/HC dimers supported by *in silico* docking and molecular dynamics (MD) simulation. Our findings clearly differentiate the cell surface dynamics and properties of empty FHCs from the peptide-loaded trimers of MHC-I, pointing to the formation of structurally well-defined HC/HC homodimers that may be responsible for distinct endosomal trafficking and other biological functions previously ascribed to FHCs.

## Results

### A micropattern assay reveals non-covalent association of MHC class I free heavy chains

In STF1 cells, which are fibroblasts that cannot load MHC-I with peptides due to a deficiency in the TAP peptide transporter, incubation at 25 °C accumulates murine HC/β_2_m heterodimers at the plasma membrane, since at that temperature, dissociation of β_2_m and subsequent endocytosis are inhibited (Day et al., 1995; Ljunggren et al., 1990; Sebastián Montealegre et al., 2015). When the temperature is shifted to 37 °C, β_2_m dissociates to yield FHCs, and lateral *in cis* interactions between HA-K^b^ and K^b^-GFP (both hybrids of the HC of the murine MHC-I molecule H-2K^b^) become visible in the micropattern two-hybrid assay, where the GFP fluorescence arranges in the shapes of the antibody micropattern (**Figure 1C**). When cognate peptide is added to the cells, HC/β_2_m/peptide trimers are stable and do not associate with each other (**Figure 2B**) (Dirscherl et al., 2018).

**Figure 2:**
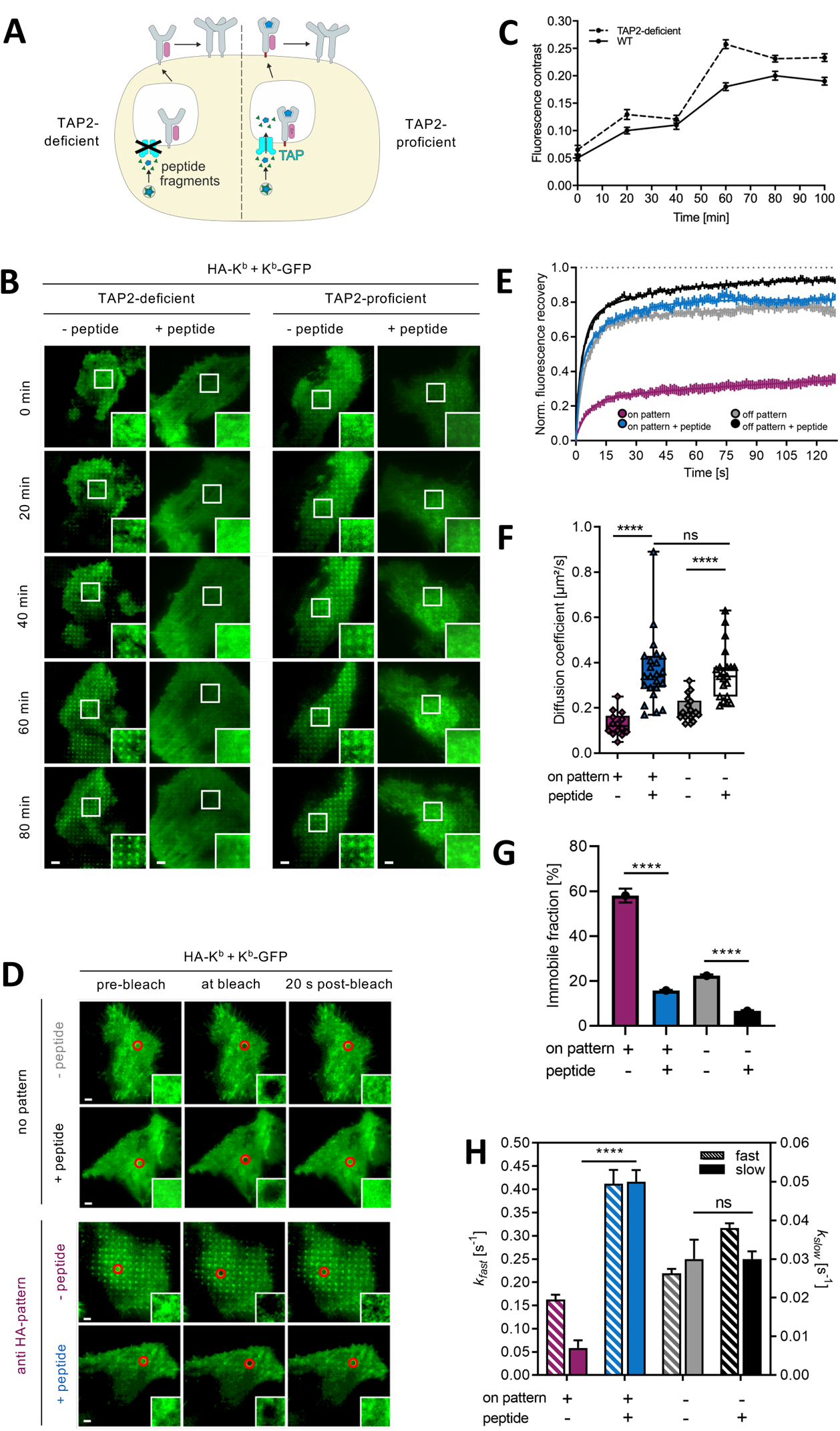
FHCs interact in TAP-proficient cells and in the absence of patterns. **A**, Schematic of cell surface interaction between FHCs in TAP2-deficient (left) and TAP2-proficient cells. **B**, Time course of the interaction between HA-K^b^ and K^b^-GFP in the anti-HA antibody micropattern assay upon temperature shift from 25 °C to 37 °C in TAP2-proficient (STF1/TAP2, right) and TAP2-deficient cells (STF1, left). Representative TIRF microscopy images are shown. 2 µM of cognate SIINFEKL peptide were added as indicated. Scale bar, 8 µm. Insets show enlarged intensity-adjusted regions within the selected cells. **C**, Quantification of fluorescence contrast from B. Error bars are standard error of the mean (SEM) of >10 cells measured in ≥2 independent experiments. **D**, FRAP analysis of K^b^ cell surface dynamics. Cells expressing K^b^ constructs as in B were monitored in the absence of patterns (top) or on anti-HA antibody patterns (bottom) with or without SIINFEKL peptide as indicated. Regions in the red circle were bleached, and recovery of fluorescence was followed over time. Scale bar, 5 μm. Insets show the enlarged intensity-adjusted region of bleached spot. **E**, Normalized mean fluorescence recovery curves from FRAP experiments, error bars represent SEM of >20 cells for each condition measured in >2 independent experiments. **F**, Diffusion coefficients calculated from E. Significant increase in diffusion coefficients upon peptide addition, both in the absence and presence of antibody patterns. ****, p ≤ 0.0001 by two-way ANOVA. ns, not significant. Error bars are SEM of >20 cells for each condition measured in ≥2 independent experiments. **G**, Immobile fraction of K^b^–GFP molecules, i.e. molecules that remain in the bleached area during the time of the experiment. With peptide vs. without peptide ****, p ≤ 0.0001 and on pattern vs. in the absence of pattern ****, p ≤ 0.0001 by two-way ANOVA. Error bars are SEM of >20 cells for each condition measured in ≥2 independent experiments. **H**, Exchange rates of K^b^ molecules calculated from fluorescence recovery curves. k_fast_ (left Y axis, hatched bars) is the rate of diffusion of K^b^–GFP molecules. k_slow_ (right Y axis, solid-color bars) is the rate of K^b^–GFP exchange on (= binding to and dissociation from) HA–K^b^. With peptide vs. without peptide ****, p ≤ 0.0001 on anti-HA pattern by two-way ANOVA. Error bars are SEM of >20 cells for each condition measured in ≥2 independent experiments.

Up to six different MHC-I allotypes are present at the cell surface of human and murine cells. To date, though, an interaction of different MHC-I allotypes in the same plasma membrane has not been shown. We therefore tested for such heterotypic interactions between K^b^ and H-2D^b^ (D^b^). Just as for K^b^/K^b^, also K^b^ and D^b^ FHCs (at 37 °C) interacted with each other, whereas β_2_m-bound heterodimers at 25 °C did not (**Figure 1D**). Hence, our micropattern assay confirmed that formation of heterotypic FHC interactions is possible.

We next asked which molecular characteristics are required for HC/HC interactions, and we first tested whether the HCs are held together by cytosolic disulfide bonds as previously suggested for certain allotypes under oxidizing conditions (Baía et al., 2016; Capps et al., 1993; Makhadiyeva et al., 2012). We replaced the single cysteine in the cytosolic tail of K^b^-GFP (residue 332) with a serine and tested for association between this mutant and wild type HA K^b^ (**Figure 1E**). K^b^(C332S)-GFP and wild type HA-K^b^ interacted in over 90% of cells, just as the wild type K^b^-GFP, demonstrating that cysteine 332 is not required for HC/HC interactions.

To test more generally for any disulfide bonding in MHC-I interactions, we immunoprecipitated HA-K^b^ molecules from STF1 cell lysate with an anti-HA antibody. By non-reducing gel electrophoresis, covalent homodimers of K^b^ were not observed, while the well-described disulfide-linked homodimers of the human MHC-I allotype HLA-B*27:05 (Dangoria et al., 2002) were readily detected (**Figure 1F**). We conclude that the K^b^ FHCs are non-covalently associated and that they do not undergo intramolecular disulfide bonding under the conditions of our assay.

Since the cysteines in the cytosolic tail of K^b^ are not required for HC/HC interaction, we hypothesized that FHCs non-covalently associate via their extracellular domains, and that the loss of β_2_m is a prerequisite for HC/HC interaction. As anticipated, a disulfide-stabilized variant of K^b^–GFP (Y84C/A139C), in which β_2_m dissociation is dramatically decreased (Hein et al., 2014), did not interact with HA–K^b^ (**Figure 1G**). In agreement with our earlier finding that covalent attachment of β_2_m to the HC also prevents HC/HC association (Dirscherl et al., 2018), this result demonstrates that HC/β_2_m heterodimers do not interact with other HC/β_2_m heterodimers, nor with FHCs. These findings demonstrate that the MHC class I complexes observed in our system are non-covalent in nature and comprise free heavy chains, devoid of β_2_m and peptide.

### Free heavy chains associate on TAP-proficient cells

So far, we used TAP2-deficient STF1 cells to obtain homogeneous populations of free heavy chains at the cell surface. Since MHC-I HC/β_2_m heterodimers and FHCs both exist at the cell surface of wild type cells (Day et al., 1995; Ljunggren et al., 1990; Ortiz-Navarrete and Hammerling, 1991, p.), we next tested whether FHCs also associate in cells with wild type TAP function (schematic in **Figure 2A**). Following the 25 °C to 37 °C temperature shift that triggers FHC formation, we performed the same anti-HA antibody two-hybrid micropatterning assay with HA-K^b^ and K^b^-GFP as above (in **Figure 1B and C**). Again, the punctate GFP fluorescence signifies the recruitment of K^b^-GFP fusion to the printed patterns of the anti-HA antibodies, mediated by the HA-K^b^ fusion. We quantified K^b^ HC/HC interaction over time in STF1/TAP2 cells, which are able to load class I molecules with peptides, by GFP fluorescence contrast analysis between pattern elements and interspaces. Interactions increased in TAP2-positive and in TAP2-deficient cells with the same dynamics and reached a maximum after ca. 60 min of incubation at 37 °C (**Figure 2B, C**), with the kinetics likely governed by the dissociation of β_2_m from the HC/β_2_m heterodimers, which occurs on this timescale (Sebastián Montealegre et al., 2015). As expected, HC/β_2_m/peptide trimers, stabilized by cognate SIINFEKL peptide, did not show any interaction (**Figure 2B**). This experiment confirms that HC/HC interactions indeed take place at the surface of TAP2-proficient cells. In addition, these experiments again confirm our earlier observations (Dirscherl et al., 2018) that the HC/HC interaction occurs only between FHCs: when dissociation of β_2_m is inhibited by low temperature (**Figure 1 DEG**) or by added peptide (**Figure 2 BD**), the interaction does not occur. The same experiments also strongly suggest that the recruitment of the K^b^-GFP fusion to the pattern elements specifically reports on the interaction between MHC class I molecules (since it depends on their conformation) and is not due to an attraction between K^b^-GFP and other surface proteins, e.g., adhesion molecules, that might have been attracted to the plasma membrane above the pattern elements.

### FHC association slows down cell surface diffusion of MHC-I

Since dimers and oligomers of FHCs have more transmembrane domains (TMDs) than single HC/β_2_m/peptide complexes, we hypothesized that they should diffuse more slowly in the plasma membrane (Gambin et al., 2006; Wilmes et al., 2015a). We therefore carried out total internal reflection fluorescence (TIRF) fluorescence recovery after photobleaching (FRAP) experiments on STF1 cells with HA-K^b^ and K^b^-GFP on surfaces with or without micropatterns (**Figure 2D**) and quantified the recovery dynamics (**Figure 2E**). Diffusion constants of FHCs (– peptide) were significantly decreased as compared to HC/β_2_m/peptide complexes (+ peptide; **Figure 2F**), suggesting that FHCs form complexes. The moderate decrease of ≈40% is in line with the formation of dimers rather than clustering into larger complexes. Since this effect occurred both in cells seeded on the pattern elements (on pattern) and in cells seeded off the micropatterns (off pattern), we conclude that the micropatterns are not required for HC/HC interactions, i.e., that the interactions are not an artefact of the two-hybrid micropattern assay. This statement is also supported by the co-immunoprecipitation of MHC-I molecules in the absence of micropatterns (Dirscherl et al., 2018; Triantafilou et al., 2000). Diffusion constants of HC/β_2_m/peptide trimers on pattern and off pattern were not significantly different, which demonstrates that the antibody micropatterns themselves do not impede the diffusion of plasma membrane proteins.

In the FRAP experiments, a portion of K^b^-GFP appeared immobile on a timescale of seconds as evidenced by the incomplete fluorescence recovery (**Fig. 2E**). This immobile fraction is significantly higher for FHCs than for HC/β_2_m/peptide trimers (**Fig. 2G**). This observation is readily explained for K^b^-GFP bound to HA-K^b^ molecules immobilized within micropatterns, but it is remarkable for the FHCs on cells outside of the micropattern (gray column in **Fig. 2G**). There, it may be ascribed to the formation of large oligomers, and/or to the association of FHCs with immobile structures, such as the cytoskeleton and/or due to sequestration in endocytic membrane compartments (Bondar et al., 2020; Ibach et al., 2015; Mylvaganam et al., 2018; Vámosi et al., 2019). From the FRAP curves, the exchange rate of the freely diffusing pool of K^b^-GFP into and out of the bleached regions of interest (ROIs) is obtained through a bi-exponential fit as the slow recovery rate (*k_slow_*) (**Figure 2H**, solid bars); the fast recovery rate *k_fast_* represents free diffusion (Sprague and Mcnally, 2005)(see the Materials and Methods). On the pattern elements, the *k_slow_* of FHCs was much smaller than in cells outside the patterns, suggesting a half-time of dissociation from the pattern-bound immobile associations of about 140 seconds. Likewise, the fluorescence signal of pattern elements in the immediate vicinity of the bleached region remained unaltered, indicating that no detectable exchange of K^b^-GFP between the enriched HA-K^b^ regions occurs on the second timescale (**Figure 2S3**). This very slow exchange of K^b^-GFP molecules associated on pattern elements suggests either multiple association and dissociation events in a small radius due to densely immobilized binding partners, or a very stable association of the immobile FHCs (see the discussion). Taken together, the data from **Figure 2** show reduced diffusion rates of K^b^ FHCs at the cell surface compared to HC/β_2_m/peptide complexes, which is most easily explained by homotypic association of the former.

### Transient FHC dimerization directly observed at single molecule level

We therefore turned to directly visualizing FHC diffusion and interaction in the plasma membrane under physiological conditions by single molecule tracking (SMT) and co-tracking (SMCT) (Moraga et al., 2015; Sevcsik et al., 2015; Wilmes et al., 2020a). STF-1 cells transiently expressing K^b^ with its N-terminus fused to monomeric GFP (GFP-K^b^) were imaged by TIRF microscopy. The GFP tag was labeled with photostable fluorescent dyes by using equal concentrations of anti-GFP nanobodies conjugated to either ATTO Rho11 (^Rho11^NB) or ATTO 643 (^ATTO643^NB), with one nanobody binding one GFP molecule, ensuring selective imaging of K^b^ in the plasma membrane and tracking with high fidelity (**Figure 3A**). Imaging was performed at 37 °C to induce formation of FHC in the absence of the peptide, while replenishment of GFP-K^b^/β_2_m dimers from the ER was inhibited by Brefeldin A (BFA).

**Figure 3:**
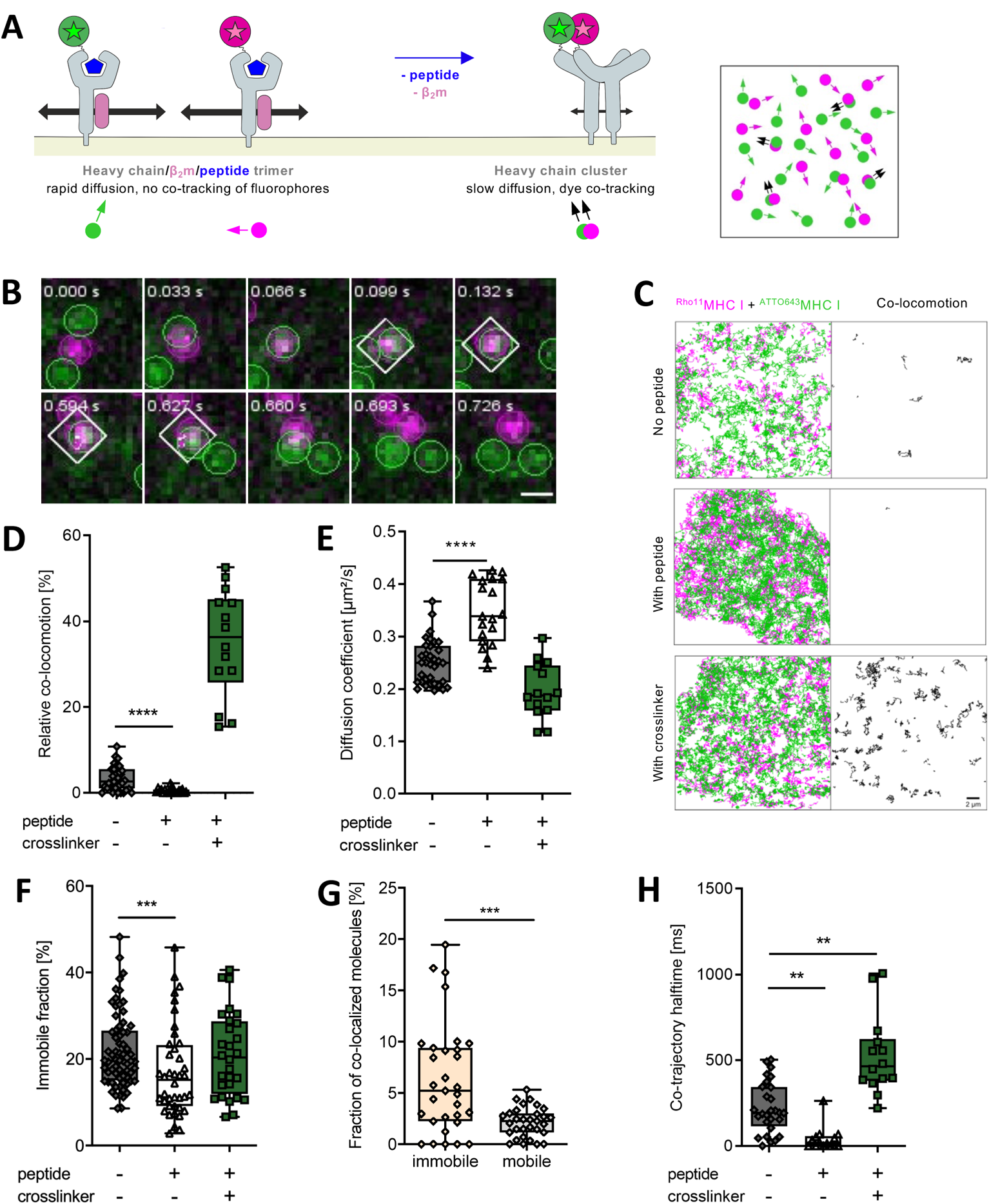
Single-molecule microscopy shows K^b^ dimer formation. **A**, Schematic on the left: analyzing FHC association by single-molecule microscopy. GFP-K^b^ molecules at the plasma membrane were labeled with two different fluorophores (ATTO Rho11, magenta, and ATTO 643, green) using anti-GFP nanobodies, with one GFP molecule binding one nanobody. Diffusion and interaction of individual molecules was quantified by tracking and co-tracking of individual molecules. Right: Top view of the plasma membrane with diffusing and co-diffusing nanobody-labeled GFP-K^b^ molecules (corresponding data are shown in **C**). **B**, Formation and dissociation of an individual GFP-K^b^ dimer detected by co-tracking analysis. Co-locomoting molecules are highlighted by a white square. Scale bar, 400 nm. **C**, Single molecule trajectories (green, magenta) and co-trajectories (black) observed for GFP-K^b^ FHCs (no peptide, top) and HC/β_2_m/peptide trimers (With peptide, middle). As a control, the same experiment was carried out after crosslinking peptide-loaded K^b^ with a tandem anti-GFP nanobody (With crosslinker, bottom). Scale bar, 2 µm. **D-H**, statistical analyses by two-sample Kolmogorov-Smirnov test (****, p ≤0.0001; ***, p ≤0.001; **, p ≤0.01). Sample numbers see the methods section. **D**, Fraction of detected co-trajectories out of the total number of observed trajectories for K^b^ in the absence and in the presence of peptide and crosslinker as indicated. **E**, Diffusion coefficients of K^b^ molecules determined by single-molecule tracking in the absence and presence of peptide and crosslinker. **F**, The fraction of immobile molecules out of the total GFP-K^b^ surface molecules determined by single-molecule tracking. SIINFEKL peptide and crosslinker were added as indicated. **G**, Comparison of the fractions of mobile and immobile GFP-K^b^ molecules that are associated. The fraction of (associated immobile K^b^)/(total immobile K^b^) is shown vs. the fraction of (associated mobile K^b^)/(total mobile K^b^). **H**, Quantification of co-trajectory half-life for GFP-K^b^ as obtained from single molecule co-trajectories with SIINFEKL peptide and crosslinker as indicated; the dissociation rates calculated from the means are 3.1 (no peptide), 17 (peptide, no crosslinker), and 1.3 s^−1^ (with peptide and crosslinker).

After labeling with ^Rho11^NB and ^ATTO643^NB, individual K^b^ subunits were observed at densities of <1 molecule/µm^2^ diffusing randomly in the plasma membrane (**Supplementary Movie S1, Supplementary Figure S3A**). Co-tracking analysis revealed homomeric interaction of K^b^ FHCs in the absence of peptide, whereas these events were very rare for peptide-loaded K^b^ (**Figure 3B-D**). As a positive control for the formation of homodimers, a crosslinker based on the tandem anti-GFP nanobody LaG16 (Fridy et al., 2014), which recognizes a different epitope than the labeled nanobodies, was added to the medium. While dimerizing of K^b^ via the GFP tag resulted in a nominal fraction of ≈38% (median) co-locomoting molecules, ≈3% of the FHCs were found associated, indicating weak interaction of FHCs at these low cell surface expression levels of K^b^ (≈1 molecule/µm^2^ in total, corresponding to <5000 molecules/cell).

From the trajectory analysis, we determined the diffusion coefficients of individual MHCI molecules by mean square displacement analysis (**Figure 3E, Supplementary Figure S3B**). In the absence of peptide, diffusion of FHCs was significantly slower than in the presence of the peptide. This decrease in single molecule diffusion coefficients supports interaction of FHCs in the plasma membrane. A similar decrease in the diffusion coefficient was observed upon dimerization of peptide-loaded K^b^ with the tandem nanobody, suggesting that FHCs associate into dimers. Indeed, largely identical diffusion coefficients were found for the fraction of molecules identified as dimers by SMCT (**Supplementary Figure S3C, Table 1**). Strikingly, the diffusion constants obtained by SMT for peptide-loaded K^b^ and for FHC from SMT are consistent with those obtained by FRAP experiments under the same conditions (**Figure 2E, Table 1**), highlighting that similar phenomena are being probed by these complementary techniques. Tracking analysis revealed a slightly elevated immobile fraction of ~20% for K^b^ in the absence of the peptide compared to ≈17% for peptide-loaded K^b^ (**Figure 3F**). Again, a similar effect was observed for artificially dimerized, peptide-loaded K^b^, corroborating the dimeric stoichiometry of associated FHCs. Importantly, the “immobile fraction” in SMT refers to much shorter time and length scales as compared to FRAP (cf. methods section), and therefore, the absolute numbers are not comparable. Within the immobile fraction, a higher level of FHC was found associated, as compared to the mobile fraction (**Figure 3G**). Overall, the single molecule diffusion analyses suggest that FHC dimers have an increased propensity to be immobilized at the plasma membrane, probably by clustering that may be related to endocytosis. Interestingly, this feature is reproduced by artificial dimerization of intact MHC-I by a crosslinker, suggesting that FHC dimerization is a switch regulating its cell surface dynamics.

**Table 1:** Comparison of diffusion coefficients (mean ± SD in µm^2^/s). NA, not applicable.

|  | **FRAP** | **SMT** |
| --- | --- | --- |
| **HC/$\beta_2m$/peptide trimers (+peptide)** | 0.35 ± 0.12 | 0.35±0.06 |
| **FHCs (–peptide)** | 0.19 ± 0.05 | 0.25±0.04 |
| **FHCs (–peptide)<sup>a</sup>** | 0.19 ± 0.05 | 0.18±0.10 |
| **Dimerized HC/$\beta_2m$/peptide trimers (+peptide, + CL)** | NA | 0.20±0.05 |
| **Dimerized HC/$\beta_2m$/peptide trimers (+peptide, + CL)<sup>a</sup>** | NA | 0.15±0.04 |

<sup>a</sup>only dimers identified by SMCT

SMCT analysis moreover revealed dissociation of FHC homomers, confirming its transient nature (**Figure 3B, Supplementary Movies S2-5**). We estimated the lifetime of FHC complexes from co-trajectory length histograms. Fitting of an exponential decay revealed a significantly shorter lifetime for FHC co-trajectories compared to the average co-tracking lifetime determined for stably NB-crosslinked K^b^, which is limited by co-tracking fidelity and photobleaching (**Figure 3H**). Taken together, the SMT and SMCT results of **Figure 3** directly confirm a transient homotypic HC/HC interaction in the plasma membrane that leads to reduced diffusion velocity and immobilization, in line with the FRAP experiments. Since the degree of association was low, and higher-order oligomerization was not observed, we propose that HC/HC interactions are weak and have dimeric stoichiometry.

### The **α**_3_ domain of K^b^ forms dimers and is sufficient for FHC association

We next explored the molecular mechanism of HC/HC interaction. Since cytosolic cysteines are not involved (**Figure 1E, F**) and dissociation of β_2_m is necessary (**Figure 1G**), we hypothesized that HC/HC interactions involve the extracellular portion of the HCs. To determine more precisely which domains are involved, we used STF1 cells in the micropattern assay that expressed constructs of K^b^ that lacked the α_1_/α_2_ domain. Remarkably, α_3_-GFP showed excellent copatterning with HA-K^b^-RFP (fused to red fluorescent protein; **Figure 4A**) and with HA-α_3_ (**Figure 4B**), demonstrating that isolated α_3_ domains can bind to each other on the surface of live cells. Indeed, the isolated, soluble α_3_ domain of K^b^, when refolded *in vitro*, showed (after testing of its folded state by fluorescence spectroscopy, **Supplementary Figure S4C**) significant formation of homodimers in size exclusion chromatography (SEC; **Figure 4C**), while no prominent higher-order complexes were observed. These α_3_ homodimers were not linked by disulfide bonds as shown by nonreducing gel electrophoresis (**Figure 4D**), just like the interaction of full-length FHC in the plasma membrane (**Figure 1F**). Given the 1:1 peak ratio observed at 10 µM protein concentration, a binding affinity of 10 to 20 µM can be estimated for this interaction. The non-covalent α_3_ homodimers were also detectable by native mass spectrometry (MS) in a concentration-dependent manner, and they easily separated into the monomers when the collision cell voltage was increased in an MS/MS experiment, suggesting a low-affinity interaction (**Figure 4E-G, Supplementary Figure S4A**). These biochemical results, together with the observations on the α_3_ constructs above, suggest that FHC dimerization in the plasma membrane is at least partly based on the intrinsic affinity between the α_3_ domains. To investigate whether the α_3_ domain is indeed required for FHC dimerization, we performed a co-patterning experiment with an HA-K^b^ fusion protein that lacked the α_3_ domain (HA-K^b^(Δα_3_)-RFP) and K^b^-GFP and found the interaction significantly reduced, but still measurable (**Supplementary Figure S4B**), which suggests that the α_1_/α_2_ domain is also involved in the interaction.

**Figure 4:**
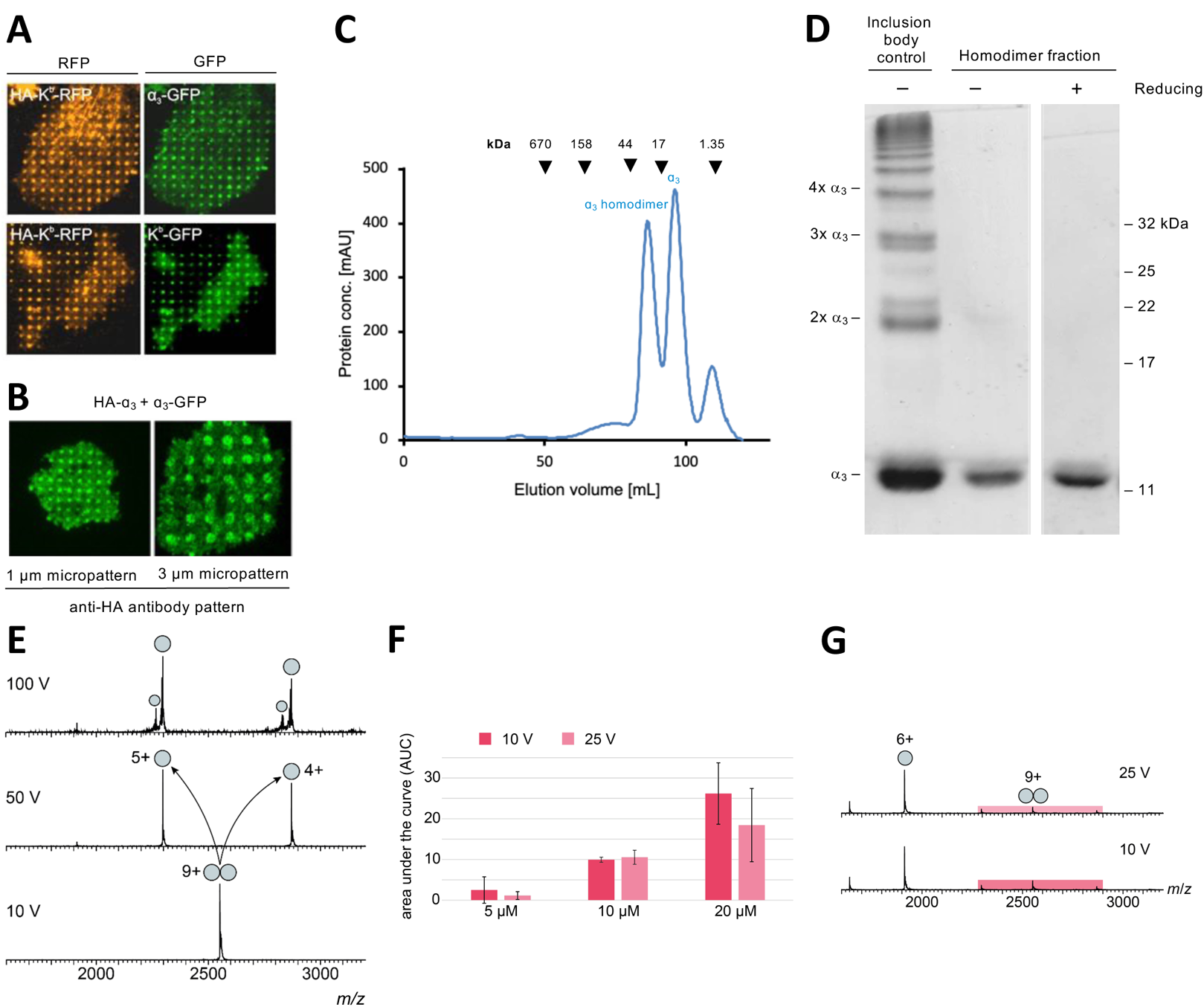
The K^b^ _3_ domains form dimers. **A**, HC/HC association between HA-K^b^-RFP and α_3_-GFP. Distance between pattern elements, 2 µm. **B**, HC/HC association between HA-α_3_ and α_3_-GFP. Distance between pattern elements, 2 µm. **C**, Size exclusion chromatography of the K^b^ α_3_ domain expressed in E.coli and folded in vitro. **D**, Nonreducing SDS-PAGE of the α_3_ domain homodimer fraction from C shows absence of disulfide linkage between the monomers. The positive control for disulfide oligomer formation is a_3_ inclusion bodies, isolated under oxidizing conditions, where some molecules are linked by disulfide bonds to form covalent oligomers as indicated on the left. **E**, MS/MS analysis of the α_3_ domain at 10 µM. The 9+ peak at 2551 *m*/*z* corresponding to the a_3_ dimer was selected for MS/MS analysis at 10 V, 50 V and 100 V in the collision cell. The dimer (double spheres) easily dissociates in the gas-phase indicating a non-covalent rather low-affinity binding event. At 50 V, no dimer is detectable anymore displaced by monomeric signal (single spheres). **F**, Overall area under the curve (AUC) for the detected α_3_ domain dimers. Native mass spectra were recorded using 5 µM, 10 µM or 20 µM α_3_ domain at 10 V (dark red) and 25 V (light red), respectively. The AUC was determined over the entire spectrum for the dimeric mass species and is shown as mean (*n*=3) ±SD. The dimeric fraction is concentration-dependent. **G**, Representive mass spectra (20 µM α_3_ domain) show the charge distributions of monomer (single sphere) and dimer (double spheres) at 10 V (dark red) and 25 V (light red) in the collision cell.

### A molecular model of the K^b^ HC dimer

To image a possible arrangement of the K^b^ heavy chains in a dimer, we used molecular docking. A starting geometry was obtained by placing the α_3_ domain of one K^b^ HC in the same position as the β_2_m in the original K^b^ HC/β_2_m heterodimer and adjusting the α_1_/α_2_ superdomain to avoid sterical overlap. The structure was then energy-minimized and further refined by Molecular Dynamics (MD) simulations (**Figure 5A**). During MD simulations of 400 ns, no signs of dissociation were observed, and a stable root-mean-square deviation (RMSD) from the start structure was reached (**Supplementary Figure S5**). In the model, the α_3_ domain of one FHC substitutes for β_2_m, binding both the α_3_ domain and the α_1_/α_2_ superdomain of another FHC with a buried interface area (BSA) of 2480 Å^2^, which is comparable to the BSA of 2740 Å^2^ between β_2_m and the HC in heterodimers (**Figure 5B**, bottom), and to other stable protein complexes (Bahadur and Zacharias, 2008). Similar results were obtained with other arrangements of the two heavy chains (not shown).

**Figure 5:**
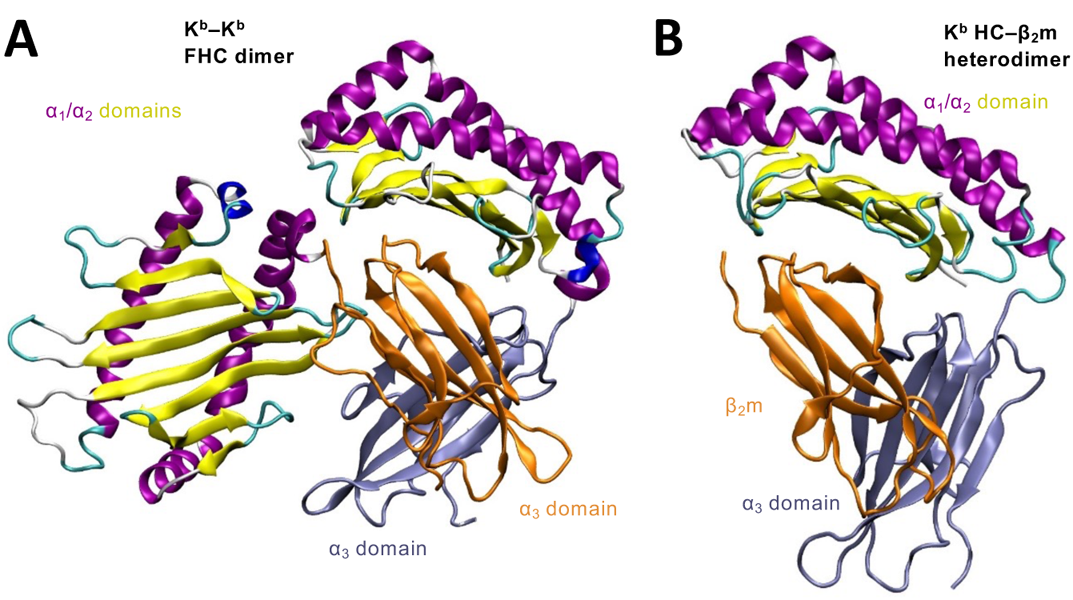
One hypothetical model of a K^b^ FHC dimer. In this hypothetical structure of the extracellular portion of an FHC dimer (**A**), one of several structures predicted by computational molecular docking and molecular dynamics simulation, the α_3_ domain of one FHC (orange) binds to another FHC (yellow/purple/lavender) in a manner resembling the binding of β_2_m in the crystal structure of peptide-loaded K^b^ (**B**, PDB 3P9L).

## Discussion

Interactions of MHC-I FHC with FHC of the same allotype, of different allotypes, and even with other cell surface proteins have been proposed to play an important role in regulating adaptive and innate immune responses (Arosa et al., 2021, 2007; Campbell et al., 2012), but the molecular principles that govern FHC interactions have remained unclear. By combining live cell interaction and diffusion analysis using cell micropatterning and FRAP as well as SMT and SMCT, we have shown here that, after losing β_2_m, murine H-2K^b^ MHC-I molecules at the cell surface interact in a homotypic and heterotypic manner to form dimers, which are transient with a stability on the second timescale. The α_3_ domains of the FHCs alone are sufficient for such interactions, but it is conceivable that the α_1_/α_2_ domain is also involved: in addition to the α_3_-α_3_ interaction, the α_3_ domain might also bind to the α_1_/α_2_ domain of the other protein, or the α_1_/α_2_ domains of the two proteins might interact with each other. We cannot currently distinguish these possibilities.

While we observed a somewhat increased tendency of FHCs to form higher oligomers, our data surprisingly identify non-covalent FHC dimers as the prevalent species at the cell surface. We analyzed the diffusion properties of FHCs and HC/β_2_m/peptide trimers at different cell surface expression levels by FRAP and SMT. FRAP experiments were carried out at high surface densities, which probably exceeded the typical endogenous cell surface expression levels of 10^5^ copies/cell (Spack and Edidin, 1986). By contrast, SMT was performed at densities of ≈1 molecule/µm^2^, *i.e.*, <5000 molecules/cell. Despite these differences in expression levels, we found highly consistent diffusion coefficients *D* by both FRAP and SMT, which revealed moderately decreased mobility upon dissociation of β_2_m (**Table 1**). However, under both conditions, the decrease in *D* was ≈40-50%, which perfectly agrees with the change in *D* upon dimerizing intact MHC-I by the tandem nanobody (**Figure 3E**). Likewise, we and others have previously observed very similar decreases of 30-50% upon dimerization of cell surface receptors (Ho et al., 2017; Low-Nam et al., 2011; Moraga et al., 2015; Richter et al., 2017; Váradi et al., 2019; Wilmes et al., 2015a, 2020a), corroborating that the mobile HC/HC complexes are mostly dimers.

Direct detection of FHC interaction by SMCT corroborated transient, non-covalent dimerization and only minor clustering into higher oligomers. Estimated lifetime of the non-covalent FHC/FHC dimer (*t_½_* = 220 ±150 ms, **Figure 3G, H**) were well within the measurable range, *i.e.*, substantially below the apparent half-life obtained for the quasi-irreversibly, nanobody-crosslinked control (*t_1/2_* >10 s), which defined the limit of co-tracking fidelity. Together with the immunoprecipitation experiments (**Figure 1F**) and micropatterning of FHC lacking free cysteines (**Figure 1E**), these observations clearly establish that the FHC molecules at the plasma membrane are not covalently linked. The short half-life and the non-covalent monomer-dimer equilibrium observed by size exclusion chromatography at higher micromolar concentration for the isolated α_3_ domain (**Figure 4CD**), as well as the mass spectrometry data (**Figure 4E-G**), point to a low-affinity interaction with a dimerization *K*_d_ of perhaps 10 to 20 µM. Previous quantitative studies on affinity-dimerization correlation of heterodimeric cytokine receptors (Wilmes et al., 2015a) would predict efficient FHC dimerization at physiological densities above 10 molecules/µm^2^, which is in line with the endogenous MHC-I expression level. The relatively low level of dimerization we observed by SMCT can be rationalized with the high background of endogenous MHC-I that are unlabeled and invisible to us. In line with this observation, the decrease of the diffusion coefficient in SMT was much more prominent than the dimer fraction identified by SMCT. This interpretation is in line with the observation that more efficient interaction was observed at the elevated cell surface expression levels used in micropatterning experiments.

The weak and transient nature of HC/HC dimerization seen by SMT is similar to the nature of cell surface protein-protein association measured in other systems (Lin et al., 2014). It does not conflict with the clear and stringent patterning of the GFP fusion in the micropattern/FRAP experiments. In the latter, the concentration of HA-tagged bait HCs is considerably higher due to their immobilization by the antibodies, which prevents their endocytosis, and this probably creates an affinity matrix for the GFP fusions that can retain them for many seconds due to rebinding events. This also suggests that in endosomes, whose internal volume is very small, HC/HC interaction might be potentiated due to the increase in the concentration of the monomers compared to the plasma membrane.

In line with the observation of non-covalent dimerization in the plasma membrane, we obtained a robust structural model of a self-contained K^b^ FHC homodimer. The atomistic model (**Figure 5**) was derived from the experimental findings that dissociation of β_2_m is required, that the α_3_ domains are sufficient, and that a direct α_3_/α_3_ interaction exists (**Figures 1E, 4A-C**). We propose that β_2_m dissociation exposes a binding site on the FHC for the α_3_ domain of another FHC. However, other arrangements of the two FHCs in a dimer are theoretically possible, and only experimental data will give a definitive answer.

Several findings in the literature are consistent with the formation of complexes of MHC class I HCs in the absence of β_2_m and peptide; this applies both to the ‘classical’, or class Ia, proteins HLA-A/B/C (and in mouse: H-2D/K/L) as well as to the ‘non-classical’, or class Ib, protein HLA-F (Armony et al., 2021; Bodnar et al., 2003; Chakrabarti et al., 1992; Matko et al., 1994; Triantafilou et al., 2000). Still, it is important to differentiate these HC/HC dimers from class I associations described elsewhere (partially reviewed in (Arosa et al., 2021, 2007; Campbell et al., 2012)), namely homo- and heterotypic HC/β_2_m/peptide trimers that are covalently dimerized via disulfide bonds in their cytosolic tails (Capps et al., 1993; Makhadiyeva et al., 2012) and that may play a role in binding the LILRB NK cell receptor (Baia et al., 2016); the non-covalent nano- and microscale clusters of HC/β_2_m/peptide trimers (not detected in our system) that may stem from the fusion of exocytic vesicles with the plasma membrane and that may play a role in TCR recognition (Blumenthal et al., 2016; Ferez et al., 2014; Fooksman et al., 2006; Lu et al., 2012); the macroscopic ‘clusters’ of class I molecules at the signaling interface between cells (Fassett et al., 2001) and the covalent dimers of the HLA-B*27:05 heavy chain that are linked by disulfide bonds through Cys-67 (Chen et al., 2017), though our non-covalent HC/HC dimers may be a precursor to the formation of the latter.

The unexpected discovery that FHCs non-covalently associate into defined dimers allows exciting hypotheses of their distinct functional properties. FHC dimers might be responsible for the immunomodulatory functions of cell surface heavy chains, *i.e.,* the stabilization of MHC-I trimers to assist T cell activation (Geng et al., 2018; Schell, 2002) and direct binding of FHCs to receptors on other cells (for example, FHCs of HLA-F binding to activating receptors on NK cells (Dulberger et al., 2017, 2017; Goodridge et al., 2013)). Furthermore, dimerization of FHCs might enhance endocytosis in order to remove the non-functional FHCs, which themselves cannot activate T cells and are known to be short-lived (Mahmutefendic et al., 2011; S. Montealegre et al., 2015); alternatively or additionally, such associated HCs may bind to other proteins *in cis* and promote their removal from the plasma membrane. Such endocytic removal might be achieved by altered endosomal routing, since the local density of membrane proteins in endosomes is higher than at the plasma membrane, and thus, efficient dimerization of MHC-I FHCs is expected. In such a scenario, even transient oligomerization in endosomes might prevent the return of internalized MHC-I FHCs to the cell surface (S. Montealegre et al., 2015). Taken together, the presence of non-covalent, transient FHC dimers points to exciting new aspects in the regulation of MHC-I functions with much potential for further investigation.

## Materials and Methods

### Cells and cell lines

TAP-deficient human STF1 fibroblasts (kindly provided by Henri de la Salle, Etablissement de Transfusion Sanguine de Strasbourg, Strasbourg, France) were cultivated at 37 °C and 5% CO_2_ in Earle’s minimum Essential Medium (MEM) with stable glutamine supplemented with 10% fetal bovine serum (FBS), non-essential amino acids and HEPES buffer without addition of antibiotics.

### Genes, vectors, and gene expression

HA-K^b^ and K^b^-GFP constructs were described previously (Dirscherl et al., 2018). HA-K^b^ carries an influenza hemagglutinin (HA) tag at the N terminus of the full-length murine H-2K^b^, whereas K^b^-GFP carries a GFP domain at the C terminus of H-2K^b^. The D^b^-GFP construct is analogous to the K^b^-GFP construct. The α_3_-GFP construct consists of the H-2K^b^ signal sequence and residues 204-369 of H-2K^b^, including transmembrane and cytosolic domains. The HA-K^b^(Δα_3_)-RFP construct, compared to HA-K^b^, lacks residues 204-294 and carries an additional RFP (red fluorescent protein) taken from pcDNA3-mRFP (see the key resources table). The GFP-K^b^ construct used in single-molecule imaging consists of a signal sequence and GFP fused to the N terminus of H-2K^b^ itself lacking a signal sequence. Amino acid numbering of K^b^ in this paragraph includes the signal sequence.

Stable cell lines were generated by lentiviral transduction as described (Hein et al., 2014), and transient transfection was achieved by electroporation (Garstka et al., 2007) or by calcium phosphate precipitation (Graham and van der Eb, 1973) as described.

### Micropattern assay

#### Photolithography

Silicon master molds were prepared by semiconductor photolithography as described previously (Dirscherl et al., 2017).

#### PDMS stamps and Antibody Patterns

PDMS stamps were generated from basic elastomer and curing agent (Sylgard 184 Silicone Elastomer Kit) as described previously (Dirscherl et al., 2017).

#### Patterning cell surface proteins

Coverslips with antibody pattern were placed into sterile 6-well plates. Cells were immediately seeded as indicated at a concentration of ca. 50 000 cells per well. Usually, cells were incubated for 4-6 hours at 37 °C for adhesion and then shifted to 25 °C to accumulate MHC-I molecules at the cell surface. Samples were then kept at 25 °C to increase cell surface heterodimer levels or shifted back to 37 °C for 3-4 hours to induce FHCs by dissociation of β_2_m.

#### Dyes

Purified antibodies were labeled with Alexa Fluor 647 NHS ester (Thermo Fisher Scientific, Darmstadt, Germany) according to the manufacturer’s protocol.

#### Peptides

The K^b^-specific peptide SL8 (SIINFEKL in the single-letter amino acid code) was synthesized by GeneCust (Ellange, Luxemburg) and emc microcollections (Tübingen, Germany) and purified by HPLC (90% purity).) Peptides were added to the cells at a final concentration of 2 µM for 15-30 min at 37 °C to induce peptide binding (Dirscherl et al., 2018).

#### Washing and fixation

Cells were washed with phosphate buffered saline (PBS), 10 mM phosphate pH 7.5, 150 mM NaCl), fixed with 3% paraformaldehyde (PFA), and observed by confocal laser scanning microscopy (cLSM).

#### Microscopy

We used a confocal laser scanning microscope (LSM 510 Meta, Carl Zeiss Jena GmbH, Germany) equipped with argon and helium-neon lasers at 488, 543, and 633 nm. Images were recorded with a 63× Plan Apochromat oil objective (numerical aperture 1.4) at a resolution of 1596 × 1596 pixels. Data acquisition was performed with the LSM 510 META software, release 3.2 (Carl Zeiss Jena). During image acquisition, patterns and cells were imaged in the same focal plane at a pinhole of 1 Airy unit. Image analysis and processing were performed using ImageJ (National Institutes of Health, Bethesda, USA). Image processing comprised cropping, rotation and adjustment of brightness and contrast levels. Experiments in Figure 1 were repeated at least three times each.

### Recombinant _3_ domain of H-2K^b^

The _3_ domain of H-2K^b^ (residues 205-295) was cloned into pET3a (preceded by the residues MAIQR and followed by DRDM) and expressed in *E. coli* BL21(*DE3*) *pLysS*, refolded *in vitro* as described, and isolated by size exclusion chromatography (SEC) on a Cytiva Hiload Superdex 200 16/600 column (Anjanappa et al., 2020). Molecular weights of the peaks were determined by comparison to SEC protein standards (Cytiva), namely bovine thyroglobulin (670 kDa), bovine gamma globulin (158 kDa), chicken ovalbumin (44 kDa), horse myoglobin (17 kDa), and vitamin B12 (1.35 kDa). The D5 fraction, corresponding to the elution peak at approximately 20-30 kDa, was boiled with or without DTT (0.6 M final concentration) in sample buffer (LSB) (350 mM Tris-Cl pH 6.8, 10.28% sodium dodecyl sulfate (SDS), 36% glycerol 0.012% bromophenol blue). Inclusion body extract boiled without DTT (non-reducing) served as positive control for the formation of covalent oligomers. Protein quality control after refolding and SEC was performed by nanoscale differential scanning fluorimetry (nanoDSF) runs (see **Figure 4S2**) acquired with a Nanotemper Prometheus NT.48 fluorimeter (Nanotemper, Munich) controlled by PR.ThermControl (version 2.1.2).

### Precipitation of surface class I

TAP2-deficient STF1 cells expressing either CE3-HA-K^b^ or CE3-HA-B*27:05 were kept overnight at 25 °C and then pretreated with tris(2-carboxyethyl)phosphine (TCEP; 1 mM, 10 min), labeled with 400 nm of Bio-MPAA-K3 (5 min) at room temperature (RT) (Reinhardt et al., 2014). Lysis was performed in native lysis buffer (50 mM Tris Cl (pH7.4), 150 mM NaCl, 5 mM EDTA, and 1% Triton X100) for 1 hour at 4 °C. Biotinylated surface proteins were then isolated with neutravidin-coated agarose beads (Thermo Fisher Scientific, Darmstadt Germany). The isolates were boiled at 95 °C for 7 min in the presence (reducing) or absence (non-reducing) of 10 mM dithiothreitol (DTT) in sample buffer as described above. Samples were separated by SDS-PAGE and transferred onto polyvinylidene fluoride (PVDF) membranes. MHC molecules were visualized on the membranes with polyclonal rabbit anti-HA antibody as primary antibody (ab9110, Abcam, Cambridge, United Kingdom) and alkaline phosphatase-conjugated anti-rabbit serum from goat as secondary antibody (1706518, Biorad, Munich, Germany). The signals were visualized by treating the blot with BCIP/NBT substrate (B1911, Sigma Aldrich, St. Louis, Missouri, United States).

### Flow cytometry

For verification of cell surface levels of MHC-I, flow cytometry was performed with anti-HLA class I antibody W6/32 (Barnstable, 1978) and anti-HA antibody (12CA5, described in key resources table). Antibody-antigen complexes were labeled with goat secondary antibody against mouse IgG conjugated with allophycocyanin (APC) (115-135-164, Dianova, Hamburg, Germany). Fluorescent signal was recorded by a CyFlow1Space flow cytometer (Sysmex-Partec, Norderstedt, Germany) and analyzed by Flowjo, LLC software.

### FRAP

**Microcontact printing and antibody patterning for total internal reflection fluorescence (TIRF) microscopy** was performed as described previously (Lanzerstorfer et al., 2020). In short, a field of a large-area PFPE elastomeric stamp (1 µM grid size), obtained by the EV-Group (St. Florian am Inn, Upper Austria, Austria), was cut out, and washed by flushing with ethanol (100%) and distilled water. After drying with nitrogen, the stamp was incubated in 50 mL bovine serum albumin (BSA) solution (1 mg/mL) for 30 min. This step was followed by washing the stamp again with PBS and distilled water. After drying with nitrogen, the stamp was placed with homogeneous pressure onto the clean epoxy-coated glass bottom of a 96-well plate and incubated overnight at 4 °C. The next day, the stamp was stripped from the glass with forceps, and the glass bottom was bonded to a 96-well plastic casting with adhesive tape (3M) and closed with an appropriate lid. For the live cell experiments, a reaction chamber was incubated with 100 µL streptavidin solution (50 µg/mL) and incubated for 30 min at room temperature. After washing two times with PBS, 100 µL biotinylated antibody solution (10 µg/mL) was added for 30 min at room temperature resulting in an antibody surface coverage of > 85% (**Supplementary Figure S1AB**). Finally, the incubation wells were washed twice with PBS, and cells were seeded at defined cell density for the live cell microscopy analysis. The cells were allowed to attach to the surface for at least 3-4 h prior to imaging to ensure a homogeneous cell membrane/substrate interface, which is a prerequisite for quantitative TIRF microscopy. Deformation of the plasma membrane on top of the pattern elements was excluded, since control cells that were stained with the lipophilic tracer DiD (1,1′-dioctadecyl-3,3,3′, 3′-tetramethylindodicarbocyanine), which stains the plasma membrane uniformly, showed no patterning (**Supplementary Figure S1C**). For negative control to test the adhesion of the antibodies, anti-HA antibody (Abcam, ab26228) was labeled with a Zenon Alexa Fluor 488 IgG Labeling Kit (Thermofisher, Z25102), printed, bleached, and fluorescence recovery was quantified as described below (**Supplementary Figure S2AB**). In a second control experiment, binding and dissociation of a construct with both tags (HA-K^b^-GFP) from the antibody micropattern was tested, which only accounted for less than 20% of the mobility of the K^b^ fraction (**Supplementary Figure S2C**). Dissociation of Kb-GFP from micropattern elements was negligible (**Supplementary Figure S2DE)**. Thus, in our experiments, *k_slow_* was determined by the binding events between K^b^-GFP and other K^b^ molecules.

#### Live-cell TIRF microscopy

The detection system was set up on an epi-fluorescence microscope (Nikon Eclipse Ti2). A multi-laser engine (Toptica Photonics, Munich, Germany) was used for selective fluorescence excitation of GFP at 488 nm and RFP at 568 nm. The samples were illuminated in total internal reflection (TIR) configuration (Nikon Ti-LAPP) using a 60x oil immersion objective (NA = 1.49, APON 60XO TIRF). After appropriate filtering with standard filter sets, the fluorescence was imaged onto a sCMOS camera (Zyla 4.2, Andor, Northern Ireland). The samples were mounted on an x-y-stage (CMR-STG-MHIX2-motorized table, Märzhäuser, Germany), and scanning of the larger areas was supported by a laser-guided automated Perfect Focus System (Nikon PFS).

#### TIR-FRAP experiments and calculation of diffusion coefficients

FRAP experiments were carried out on an epi-fluorescence microscope as described above. Single patterns (or equivalent ROIs with 1 µM in diameter in unpatterned cells) were photobleached (Andor FRAPPA) with a high-intensity laser pulse (488 nm) applied for 500 ms. Recovery images were recorded at indicated time intervals. Normalization of data was done by pre-bleach images, and first data analysis was carried out using NIS Elements software package (Nikon). Further data processing was done in Graphpad Prism. Resulting FRAP curves were plotted based on the standard error of the mean and fitted using a bi-exponential equation. Kinetic FRAP parameters were directly obtained from curve fitting with the following model assumptions: Since unhindered MHC-I diffusion (e.g., off pattern and/or after peptide treatment; Fig. 2E) was found to be much faster than the recovery within our pattern elements, we applied a diffusion-uncoupled recovery scheme (Sprague and Mcnally, 2005). Here, after the photobleaching step, fluorescent molecules rapidly diffuse throughout the bleached micropattern, and only bound bleached molecules remain inside the spot. The bleached molecules then gradually dissociate from their binding sites. Unbleached molecules can then replace the bleached molecules at the binding sites as they become vacant. The diffusion-uncoupled FRAP recovery curve consists of two separable components: the early recovery due to diffusion and the slower recovery due to exchange at binding sites, and can be fitted using a diffusion-uncoupled two-component fit:

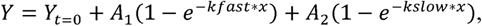

where A_1_ is the amplitude of the fast-diffusing population, A_2_ the amplitude of the slow diffusing population (binding reaction), and k_fast_ and k_slow_ are the rate constants of A_1_ and A_2_, respectively.

Diffusion coefficients were obtained using the initial image recordings and the simFRAP plugin for ImageJ (Blumenthal *et al*., 2015).

#### Temperature-induced K^b^ associations

Temperature-dependent experiments were carried out on an epi-fluorescence microscope as described above further equipped with a cage incubator (Okolab, Shanghai, China). Cells were grown at 25 °C overnight on antibody-patterned surfaces and treated with SIINFEKL peptide as indicated. For induction of K^b^ FHC association, cells were mounted on pre-warmed microscopy stage, and imaging of the GFP signal was started when the medium reached 37 °C.

#### Fluorescence contrast quantitation

Contrast analysis was performed as described previously (Lanzerstorfer et al., 2014) In short, initial imaging recording was supported by the Nikon NIS Elements software. Images were exported as TIFF frames and fluorescence contrast analysis was performed with the Spotty framework (Borgmann et al., 2012). The fluorescence contrast <c> was calculated as

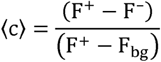

where F^+^ is the intensity of the inner pixels of the pattern, F^−^ the intensity of the surrounding pixels of the micropattern, and F_bg_ the intensity of the global background.

### Single molecule microscopy

#### Imaging

Cells were transferred 48 h post transfection onto glass coverslips coated with a poly-L-lysine-graft-(polyethylene glycol) copolymer functionalized with RGD tripeptide to minimize non-specific binding of fluorescent nanobodies (You et al., 2014). Cells imaged in presence of SL8 peptide (sequence SIINFEKL) were pre-incubated with 1 µM of SL8 peptide 12 h before imaging. Single-molecule imaging experiments were conducted by total internal reflection fluorescence (TIRF) microscopy with an inverted microscope (Olympus IX83) equipped with a motorized 4-Line TIR illumination condenser (Olympus) and a back-illuminated electron multiplying (EM) CCD camera (iXon Ultra 897, Andor Technology). A 100× magnification objective with a numerical aperture of 1.45 (UPLAPO 100× HR, NA 1.5, Olympus) together with a 1.6 × magnification changer was used for TIR illumination of the sample. Imaging was conducted with or without 1 µM of fresh SL8 peptide. The sample was preincubated with 10 µg/mL of Brefeldin A (BFA) for 15 min in order to inhibit protein transport of GFP-K^b^/β_2_m heterodimers to the plasma membrane. All experiments were carried out at 37 °C in medium without phenol red supplemented with an oxygen scavenger and a redox-active photoprotectant to minimize photobleaching (Vogelsang et al., 2008) and penicillin and streptomycin (PAA). For cell surface labeling of GFP-K^b^, a 1:1 mixture of anti-GFP nanobodies (2 nM each) site-specifically conjugated with ATTO 643 and ATTO Rho11 (Wilmes et al., 2020b), respectively, were added to the medium, thus ensuring >90 % binding given the 0.3 nM binding affinity (Kirchhofer et al., 2010). After incubation for at least 5 min, image acquisition was started with the labeled nanobodies kept in the bulk solution during the whole experiment in order to ensure high equilibrium binding. Dimerization of the positive control was induced by applying 0.3 nM of a tandem nanobody crosslinker (LaG16V2) binding to an orthogonal epitope (Fridy et al., 2014). For single molecule colocalization and co-tracking experiments, orange (ATTO Rho11) and red (ATTO 643) emitting fluorophores were simultaneously excited by illumination with a 561 nm laser (MPB Communications) and a 642 nm laser (MPB Communications). Fluorescence was detected with a spectral image splitter (QuadView QV2, Photometrics) with a dichroic beam splitter (Chroma) combined with the bandpass filter 600/37 (BrightLine HC) for detection of ATTO Rho11 and 685/40 (Brightline HC) for detection of ATTO 643 dividing each emission channel into 256 × 256 pixels. Image stacks of 150 frames were recorded for each cell at a time resolution of 32 ms/frame. Diffusion constants were determined by mean square displacement analysis within a time window of 320 ms (10 frames).

#### Single molecule analysis

MHC homodimerization was quantified based on sequential co-localization and co-tracking analysis using self-written Matlab code “SlimFAST “ as described in detail previously (Roder et al., 2014; Wilmes et al., 2015b). After aligning ATTO 643 and ATTO Rho11 channels with sub-pixel precision through spatial transformation based on a calibration measurement with multicolor fluorescent beads (TetraSpeck microspheres 0.1 μm, Invitrogen), individual molecules detected in both spectral channels of the same frame within a distance threshold of 100 nm were considered to be co-localized. For SMCT analysis, the multiple-target tracing (MTT) algorithm was applied to this dataset of co-localized molecules to reconstruct co-locomotion trajectories (co-trajectories) from the identified population of co-localizations. For the co-tracking analysis, only co-trajectories with a minimum of 10 consecutive steps (320 ms) were considered. This cut-off was determined based on systematic analysis of a negative control experiment with non-interacting model transmembrane proteins (Wilmes et al., 2020b) in order to minimize background from random co-localization. The relative fraction of co-tracked molecules was determined with respect to the absolute number of trajectories from both channels and corrected for homodimers stochastically double-labeled with the same fluorophore species as follows:

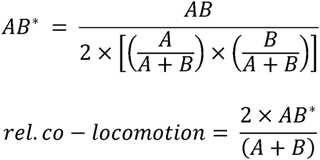

where A, B, AB and AB* are the numbers of trajectories observed for ATTO Rho11, ATTO 643, co-trajectories and corrected co-trajectories, respectively.

For **Figures 3F and 3G**, the relative fraction of co-localized mobile and immobile molecules was determined with respect to the absolute number of mobile or immobile molecules from both channels and corrected for homodimers stochastically double-labeled with the same fluorophore species as follows:

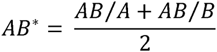

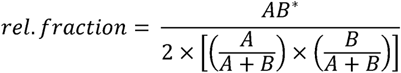

where A, B, AB and AB* are the numbers of molecules part of a trajectory observed for ATTO Rho11, ATTO 643, co-localized molecules and corrected co-localized molecules, respectively.

Immobile molecules were classified by their appearance within a radius described by the localization precision and molecular observation probability.

#### Statistics and statistical analysis

For **Figure 3D-H**, each data point represents the analysis from one cell, with ≥14 cells measured per experiment and many trajectories analyzed per cell (**3D**, 238-2513; **3E**, 238-2513; **3F**, 314-2732; **3G**, 41-354 tracked immobile particles and 362-3141 mobile particles; **3H**, 2-301 co-trajectories). Statistical significances were calculated by two-sample Kolmogorov-Smirnov test, Kruskal-Wallis test with multiple comparisons, unpaired *t* test and two-way ANOVA with multiple comparisons as indicated in the figure legends, using version Prism 8.4.0 for MacOS (GraphPad, San Diego, USA). Box plots were used for visualization and indicate the data distribution of second and third quartile (box), median (line), mean (square) and data range (whiskers).

### Native mass spectrometry

In advance of native MS measurements, a small amount (<1%) of covalent dimers of the α_3_ domain, which had formed as side product during refolding, were removed by size exclusion chromatography on Superdex 75 10/300 GL (Cytiva). Amicon Ultra 0.5 mL centrifugal filter units (molecular weight cut-off 3 kDa; Merck Millipore) were used at 14,000 x g and 4 °C to exchange purified protein samples to 150 mM ammonium acetate (99.99 %; Sigma-Aldrich), pH 7.2. The final concentration of the α_3_ domain monomer was 5 µM, 10 μ or 20 µM. Native MS analysis was implemented on a Q-Tof II mass spectrometer in positive electrospray ionization mode. The instrument was modified to enable high mass experiments (Waters and MS Vision, (van den Heuvel et al., 2006)). Sample ions were introduced into the vacuum using homemade capillaries via a nano-electrospray ionization source in positive ion mode (source pressure: 10 mbar). Borosilicate glass tubes (inner diameter: 0.68 mm, outer diameter: 1.2 mm; World Precision Instruments) were pulled into closed capillaries in a two-step program using a squared box filament (2.5 mm × 2.5 mm) within a micropipette puller (P-1000, Sutter Instruments). The capillaries were then gold-coated using a sputter coater (5.0 × 10^−2^ mbar, 30.0 mA, 100 s, 3 runs to vacuum limit 3.0 × 10^−2^ mbar argon, distance of plate holder: 5 cm; CCU-010, safematic). Capillaries were opened directly on the sample cone of the mass spectrometer. In regular MS mode, spectra were recorded at a capillary voltage of 1.45 kV and a cone voltage of 100 V to 150 V. Protein species with quaternary structure were assigned by MS/MS analysis. These experiments were carried out using argon as collision gas (1.2 × 10^−2^ mbar). The acceleration voltage ranged from 10 V to 100 V. Comparability of results was ensured as MS quadrupole profiles and pusher settings were kept constant in all measurements. A spectrum of cesium iodide (25 g/L) was recorded on the same day of the particular measurement to calibrate the data.

All spectra were evaluated regarding experimental mass (MassLynx V4.1, Waters) and area under the curve (AUC; UniDec (Marty et al., 2015)) of the detected mass species. The values of the shown averaged masses and AUC of the different species as well as the corresponding standard deviation result from at least three independent measurements. Exact experimental masses are presented in **Supplementary Figure S4A**.

### Molecular model of the K^b^ heavy chain dimer

The crystal structure of K^b^ in complex with a chicken ovalbumin epitope (PDB 3p9m) served as template to generate a model of the heavy chain dimer. Using the Pymol program (DeLano, 2002), it was possible to superimpose a copy of the 3p9m heavy chain on the β_2_m subunit of a second 3p9m structure (FHC dimer). The superposition involved only the matching of the α_3_ domain backbone of the K^b^ molecule (residues 181-277) onto the β_2_m subunit resulting in a small root-mean-square deviation (RMSD) of 1.25 Å. Very little steric overlap of the superimposed α_3_ domain with the second heavy chain was detected. This minimal overlap of the α_1_/α_2_ domain of the superimposed heavy chain was removed by adjusting backbone dihedral angles of residues 179-181 in the linker between α_1_/α_2_ domain and the α_3_ domain. The initial structural model was energy-minimized to remove any residual steric overlap and was prepared for molecular dynamics (MD) simulations using the Amber18 package (Case et al., 2018).

### Molecular docking

For comparison of FHC homodimers and K^b^ HC/β_2_m heterodimers, MD simulations were performed starting from the coordinates of the K^b^ structure in PDB 3p9m. Proteins were solvated in octahedral boxes including explicit sodium and chloride ions (0.15 M) and explicit TIP3P water molecules keeping a minimum distance of 10 Å between protein atoms and box boundaries (Jorgensen et al., 1983). The parm14SB force field was used for the proteins and peptides (Maier et al., 2015). The simulation systems were again energy minimized (5000 steps) after solvation followed by heating up to 300 K in steps of 100 K with position restraints on all heavy atoms of the proteins. Subsequently, positional restraints were gradually removed from an initial 25 kcal·mol^−1^·Å^−2^ to 0.5 kcal·mol^−1^·Å^−2^ within 0.5 ns followed by a 1 ns unrestrained equilibration at 300 K. All production simulations were performed at a temperature of 300 K and a pressure of 1 bar. The hydrogen mass repartition option of Amber was used to allow a time step of 4 fs (Hopkins et al., 2015). Unrestrained production simulations for up to 400 ns were performed. The interface packing was analyzed by calculation of the buried surface area using the Shrake method, (Shrake and Rupley, 1973); analysis of root-mean-square deviations (RMSD) was performed using the cpptraj module of the Amber18 package.

### High-speed atomic force measurements for antibody density estimation

We employed high-speed atomic force microscopy (HS-AFM), capable of resolving individual antibodies on surfaces (Preiner et al., 2014; Strasser et al., 2020) to estimate the antibody surface density in our pattern elements. Antibody micropatterns were prepared as described above. Antibody solution (10 μg/mL) was finally incubated for 15 – 1200 s, followed by washing and imaging in PBS. HS-AFM (RIBM, Japan) was operated in tapping mode at room temperature with free amplitudes of 1.5-2.5 nm and an amplitude set point of larger than 90%. Silicon nitride cantilevers (USC-F1.2-k0.15, Nanoworld AG, Neuchâtel, Switzerland) with nominal spring constants of 0.1-0.15 N/m, resonance frequencies of ~ 500 kHz,and a quality factor of ~2 in liquids were used.

Since the antibody coverage after 20 min incubation was too high to discriminate and count individual molecules, the images of the shortest incubation time (15 s) were used and thus lowest coverage and largest distance between individual molecules to estimate the area coverage per antibody (neglecting potential tip convolution effects). A height threshold of 2.7 nm above the streptavidin layer was applied for detection of particle density and mean area per particle (~190 nm ^2^), which were then used to convert the total area above the threshold in the other images (30 s – 20 min) into a surface density. The resulting surface density _Γ_ of antibodies as a function of incubation time is reasonably described by a mono-exponential function as typical for a pseudo-first order reaction (Artelsmair et al., 2008), i.e.

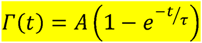

with A = 7646 ± 1828 IgGs/μm^2^ (and *τ* = 77 ± 43 s). Image analysis was performed using threshold algorithms in Gwyddion (Nečas and Klapetek, 2012).

## Acknowledgements

We thank the donors of reagents as mentioned in the Materials, Venkat Raman Ramnarayan for comments on the manuscript, Christian P. Richter for support with SMT/SMCT evaluation, Ankur Saikia and Christian Guenther for performing protein chromatography, the iBiOs staff for technical support with single molecule microscopy, the SPC facility at EMBL Hamburg for technical support, and Uschi Wellbrock for excellent technical assistance.

## Funding

Deutsche Forschungsgemeinschaft (DFG SP583/7-2), Bundesministerium für Bildung und Forschung (BMBF, 031A153A); Tönjes Vagt Foundation (XXXII), iNEXT-Discovery (11911), Jacobs University (all to SSp); Deutsche Forschungsgemeinschaft (SFB 944, projects P8 and Z, Facility iBiOs, PI 405/14-1) to JP. PL and JW acknowledge funding from the province of Upper Austria as a part of the FH Upper Austria Center of Excellence for Technological Innovation in Medicine (TIMed Center) and the Christian Doppler Forschungsgesellschaft (Josef Ressel Center for Phytogenic Drug Research). CU acknowledges funding from the Leibniz Association through SAW-2014-HPI-4 grant. The Heinrich-Pette-Institute, Leibniz Institute for Experimental Virology is supported by the Free and Hanseatic City Hamburg and the German Federal Ministry of Health.

## Author Contributions

The experimental work was performed as follows: CD, Fig. 1CDEG; ZH, Fig. 1F; PL, Fig. 2, 4AB, S1, S2, S4B; SL, Fig. 3, S3, supplementary movies; ARH, Fig. 4CD and S4B; JDK, Fig. 4E-G, S4A; MZ, Fig. 5, S5; AK and JPr, Figure S1AB. MGA, CU, JW and JPi supervised and interpreted experiments. MZ performed all MD simulations (Fig. 5, 5S1). SSp conceived the work, supervised and coordinated experiments, and wrote the manuscript in cooperation with JPi and PL and assistance by NL with input from all authors. Funding was acquired by CU, JPi, JW, MGA, MZ, PL, and SSp.

## Competing Interests

The authors declare that no competing interests exist.

## Notes

### Competing Interest Statement

The authors have declared no competing interest.

