## Supplement for "Dissociation of β_2_m from MHC Class I Triggers Formation of Noncovalent, Transient Heavy Chain Dimers"

### Supplementary Data

### Key Resources Table

| Reagent type (species) or resource | Designation | Source or reference | Identifiers | Additional Information |
|---|---|---|---|---|
| gene (Mus (M.) musculus) | H-2$K^b$ | NCBI GenBank | NM_001001892.2 | Used for plasmid design |
| gene (M. musculus) | H-2$D^b$ | NCBI GenBank | NM_010380.3 | Used for plasmid design |
| gene (M. musculus) | TAP2 | NCBI GenBank | NM_011530.3 | Used for plasmid design |
| strain, strain background (Escherichia (E.) coli Rosetta) | BL21(DE3) pLysS | Novagen | Cat. # 70956 | Used for expression of recombinant $a_3$ domain of $K^b$ |
| cell line (homo sapiens) | STF1 | PMID:10074495 | N/A | Used for all experiments |
| recombinant DNA reagent | pET3a (plasmid) | Novagen | Cat. # 69418-3 | Used for transformation of E. coli |
| recombinant DNA reagent | puc2CL6IPwo (lentiviral vector) | PMID: 21248040 | N/A | Used for transduction and transfection |
| recombinant DNA reagent (M. musculus) | puc2CL6IPwo/ E3-HA-$K^b$ | DOI: 10.7554/eLife.34150.001 | N/A | For stable transduction of STF1 cells |
| recombinant DNA reagent (M. musculus) | puc2CL6IPwo/ $K^b$-GFP | DOI: 10.7554/eLife.34150.001 | N/A | For stable co-transduction of HA-$K^b$ expressing STF1 cells |
| recombinant DNA reagent (M. musculus) | puc2CL6IPwo/ $D^b$-GFP | DOI: 10.7554/eLife.34150.001 | N/A | |
| recombinant DNA reagent (M. musculus) | puc2CL6IPwo/ TAP2 | This paper | N/A | |
| recombinant DNA reagent (M. musculus) | puc2CL6IPwo/$K^b$(Y84C/ A139C)-GFP (plasmid) | DOI: 10.1242/jcs.145334 | N/A | 10 µg for transfection of one 10 cm plate of confluent cells |
| recombinant DNA reagent (M. musculus) | puc2CL6IPwo/$K^b$(C332S) -GFP | This paper | N/A | |
| recombinant DNA reagent (M. musculus) | puc2CL6IPwo/$a_3$ domai n (of $K^b$)-GFP | This paper | N/A | |
| recombinant DNA reagent (M. musculus) | puc2CL6IPwo/ HA-$K^b$($\Delta a_3$)-GFP | This paper. RFP sequence taken from https://www.addgene.org/13032/ | N/A | |
| antibody | 12CA5 (anti-hemagglutinin (HA)) | PMID: 6192445. (Niman et al., 1983)[7] | N/A | Produced and purified in house from hybridoma cells 0.6 mg/mL for printing; 1:100 dilution of hybridoma supernatant for Western blotting. |
| antibody | polyclonal rabbit anti-HA | Abcam | Cat. # ab9110 | Used as primary antibody in Western blot, according to manufacturer's recommendations |
| antibody | alkaline phosphatase-conjugated goat anti-rabbit serum | Biorad | Cat. #1706518 | Used as primary antibody in Western blot, according to manufacturer's recommendations |
| peptide, recombinant protein | SIINFEKL peptide | GeneCust Ellange, Luxemburg | N/A | 2 mM final concentration |
| commercial assay or kit | HiLoad® 16/600 Superdex® 200 pg (column for SEC) | GE Healthcare | Cat. # GE28-9893-35 | Used for purification of recombinant $a_3$ domain of $K^b$ |
| chemical compound, drug | Alexa Fluor 647 NHS ester | Thermo Fisher Scientific | Cat. # A37566 | Used for labelling anti-HA antibodies for micropattern assay according to the manufacturer's protocol. |
| chemical compound, drug | Brefeldin A Solution (1,000X) | BioLegend | Cat. # 420601 | 10 µg/mL |
| software, algorithm | ImageJ | National Institutes of Health | https://imagej.nih.gov/ij/ | Used for image processing (cropping, rotation and adjustment of brightness and contrast levels) |
| software, algorithm | GraphPad Prism version 8.4.0 | GraphPad Software | https://www.graphpad.com | |

#### Supplementary Figure S1

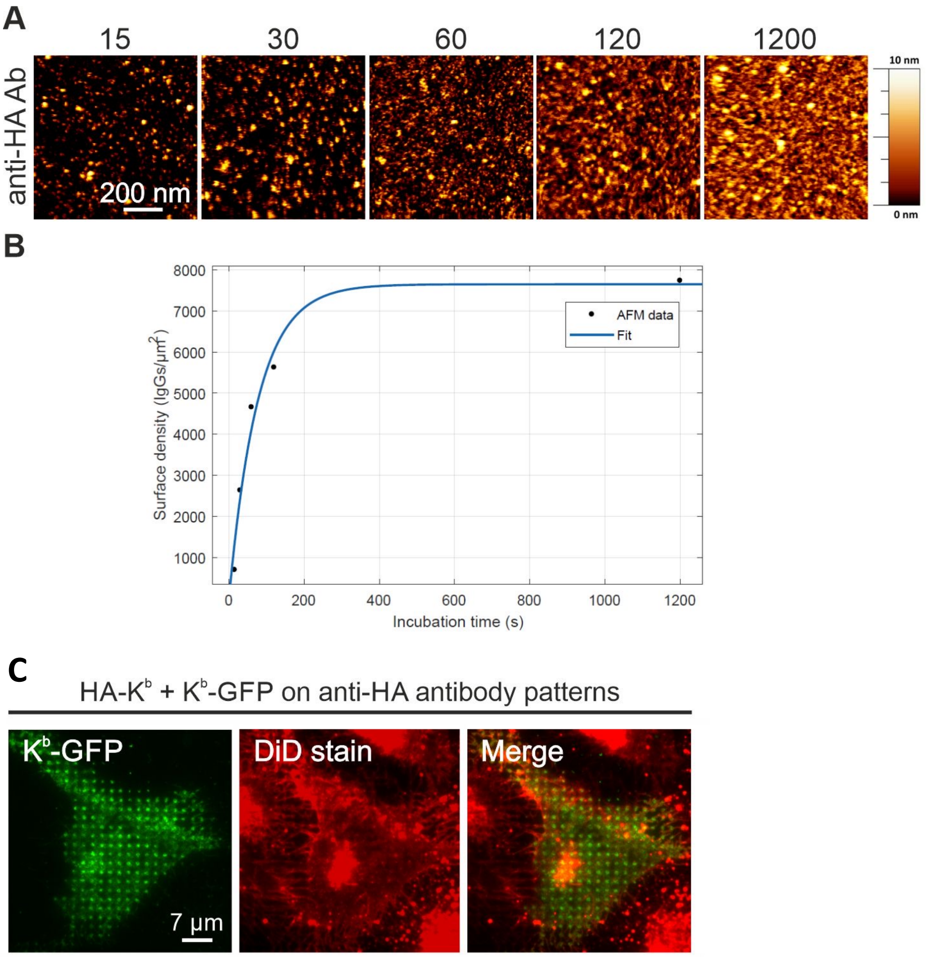


**Estimation of antibody surface densities in pattern elements by HS-AFM. (A)** anti-HA antibodies were incubated on streptavidin-coated mica sample disks (50 µg/ml) and subsequently incubated with 10 µg/ml antibody for indicated time periods (sec) followed by HS-AFM imaging. **(B)** Based on respective threshold images, the antibody surface density for each time point was calculated.

**DiD membrane stain to visualize membrane-substrate interface. (C)** STF1 cells expressing HA-Kb and Kb-GFP were grown on anti-HA antibody patterned surfaces and cell membrane was uniformly labelled by the lipophilic tracer DiD. Homogenous signal confirms sufficient cell attachement of the cells to the functionalized surface.

### Supplementary Figure S2

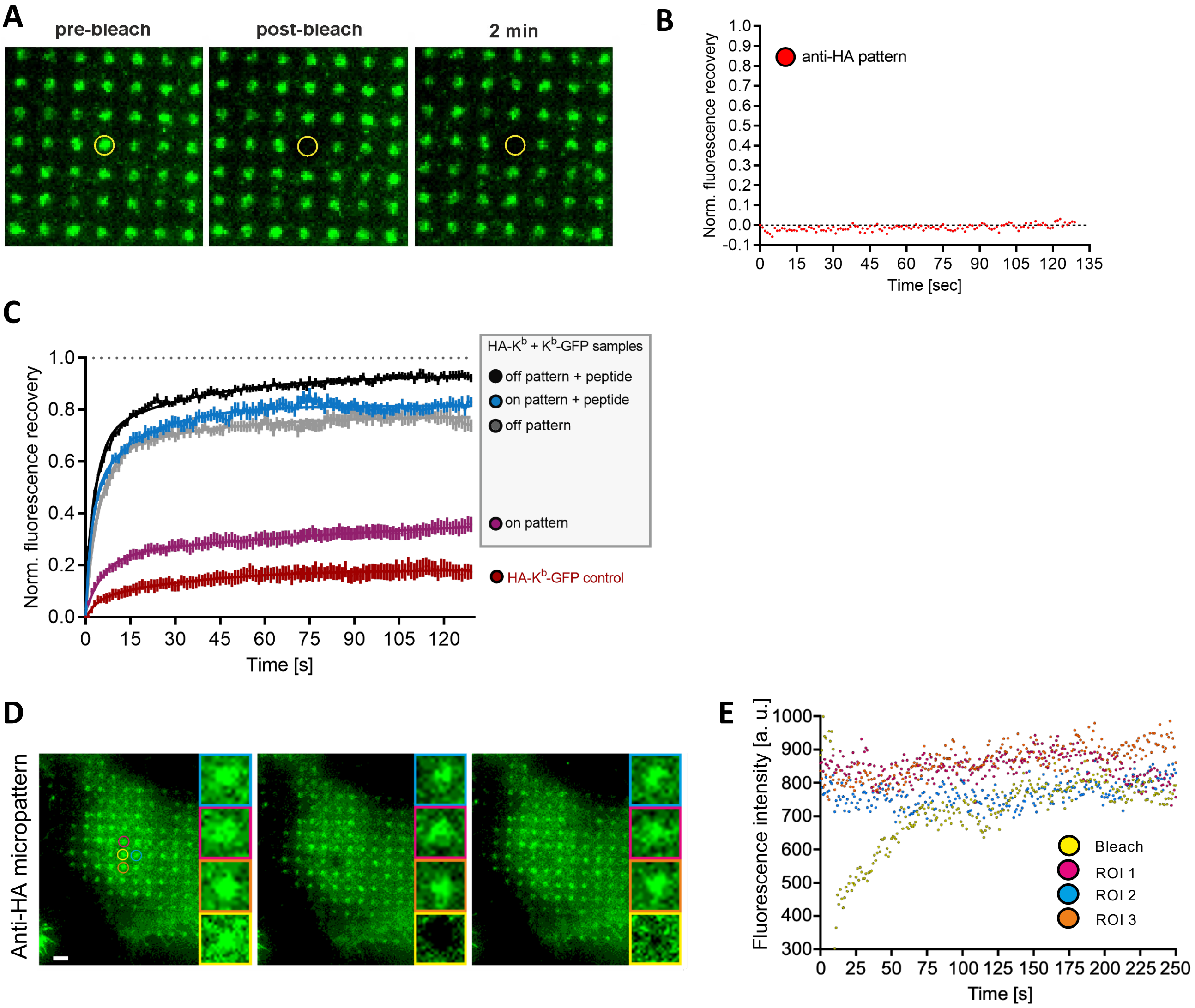


**Supplementary Figure S2:**

**Micropattern negative antibody control.** The fluorescent signal of the bleached area in an Alexa Fluor 488-labeled anti-HA antibody micropattern remains low. A pattern element was bleached (**A**), and fluorescence recovery was quantified over a time of 2 min (**B**). Pattern element distance, 2 µm.

**Binding and dissociation of HA-K$^b$-GFP to and from antibody micropattern is negligible. (C)** STF1 cells expressing HA-K$^b$-GFP (a construct with dual tags: HA at the N terminus, and GFP at the C terminus) were seeded onto anti-HA antibody patterns, GFP was bleached over one pattern element, and fluorescence recovery was measured over time (red curve). The data sets of HA-K$^b$ + K$^b$-GFP samples of Figure 2E with or without peptide are also shown as a comparison. Error bars represent SEM (*n* = 9 cells for HA-K$^b$-GFP cells).

**Dissociation of K$^b$-GFP from micropattern elements is negligible. (D)** GFP bleaching over one pattern element (yellow frame) does not affect the fluorescence of neighboring pattern elements (blue, red, orange frames) over a time period of 250 s. **(E)** quantification.

#### Supplementary Figure S3

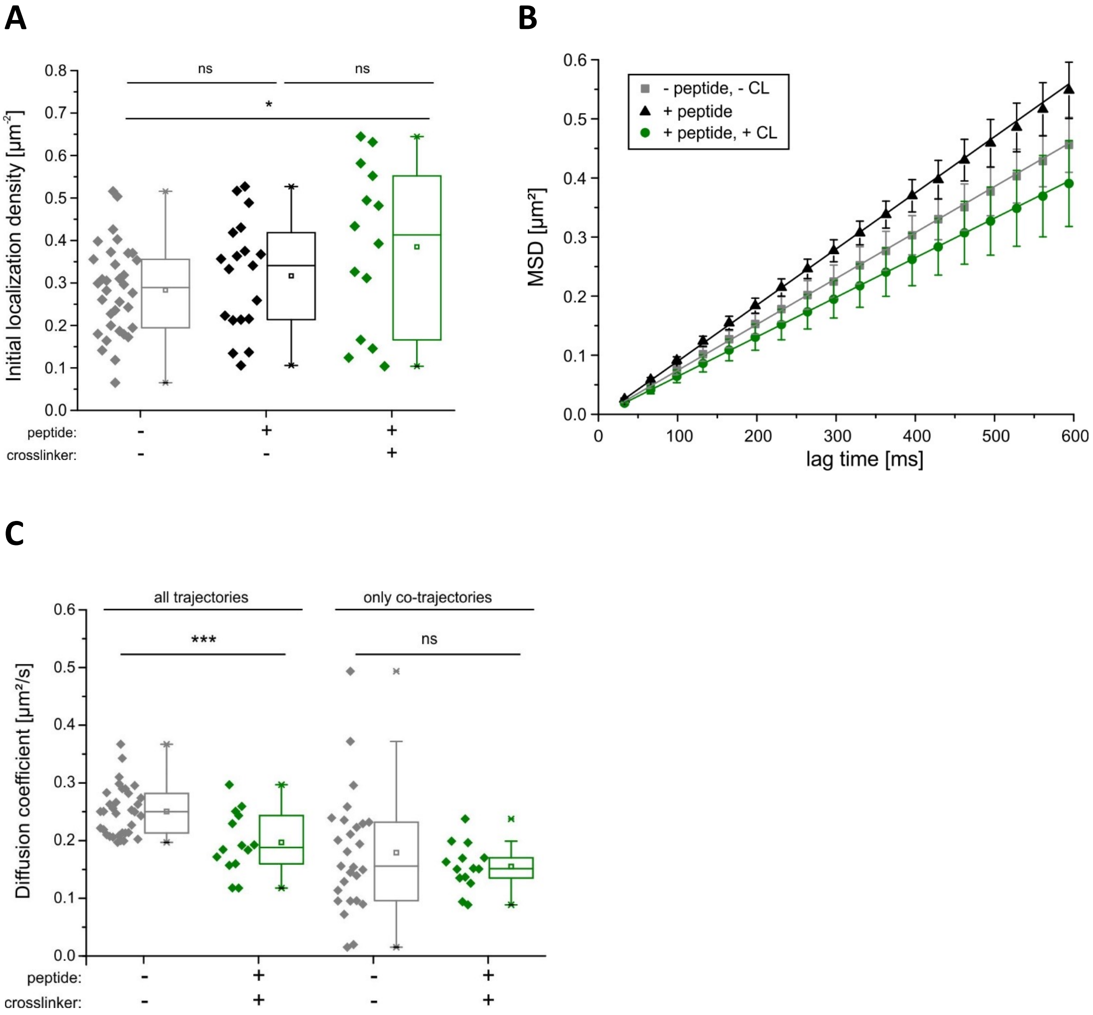


**(A) Density of molecules in SMT and SMCT experiments.** Mean initial density of localized GFP-$K^b$ molecules in each detection channel for the different conditions. Each data point corresponds to the analysis from one single cell with ≥14 cells measured for each con-dition. *, $p \le 0.05$ by a two-sample Kolmogorov-Smirnov test.

**(B) Random diffusion of mobile fraction.** Mean squared displacement (MSD) analysis of GFP-$K^b$ molecules under different conditions and linear fit. Mean of the MSD values obtained from 10 cells, error bars indicate standard deviations (SD).

**(C) Similar diffusion coefficient of FHC and MHC I dimers.**
Comparison of the diffusion constants of the entire mobile fractions and the dimers only for of GFP-$K^b$ molecules in the absence of peptide and in the presence of peptide and the tandem nanobody crosslinker (CL). ***, $p \le 0.001$ by a two-sample Kolmogorov-Smirnov test.

#### Supplementary Figure S4

**A**

| mass species | $M$ [Da] | $M_{exp}$ [Da] ± SD |
|---|---|---|
| $\alpha_3$ monomer | 11,475 | 11,472.7 ± 0.4 |
| $\alpha_3$ monomer$_{short}$ | 10,623 | 10,0621 ± 1 |
| $\alpha_3$ dimer | 22,950 | 22,947 ± 1 |
| $\alpha_3$ dimer$_{short}$ | 22,098 | 22,094 ± 1 |

**B**

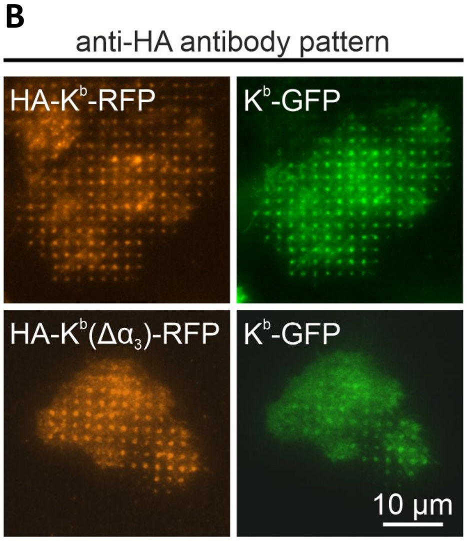


**C**

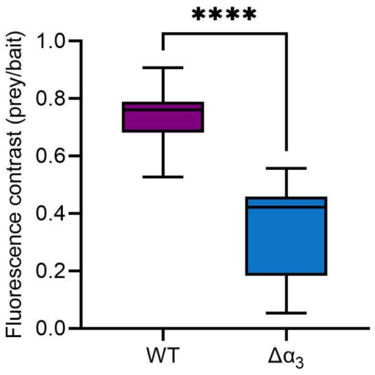


**D**

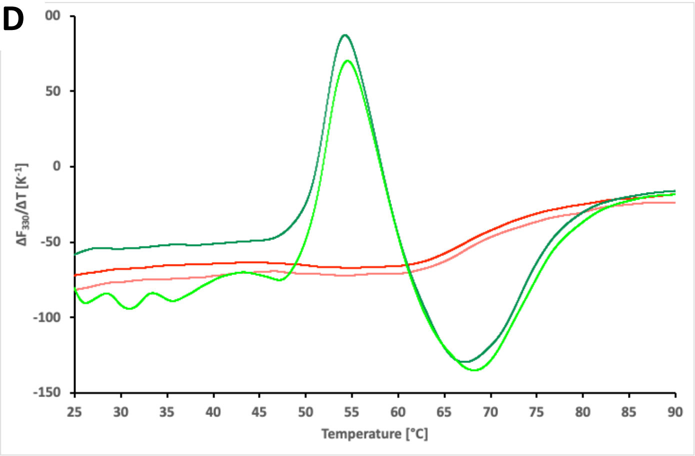


**(A) Results from native mass spectrometry.** Experimental masses ($m_{exp}$) of the different $\alpha_3$ domain protein species were determined from at least three independent mass spectrometry measurements. They are listed together with the respective values for standard deviation (SD) along with the theoretically calculated molecular weight (*M*).

**Impact of $\alpha_3$ domain on FHC dimerization.** The $\alpha_3$ domain of H-2K$^b$ was truncated and ability of FHC dimerization was compared to WT H-2K$^b$. (**B**) Representative TIRF microscopy images of STF1 cells transiently transfected with HA- Kb-RFP (WT or $\Delta\alpha_3$-mutant) and H-2K$^b$-GFP and grown on anti-HA micropatterns. (**C**) Boxplots show quantitation of bait-normalized prey fluorescence contrast of at least 10 analyzed cells. ****, $p < 0.001$ by unpaired t-test.

**(D) Proof of folding of the recombinant $\alpha_3$ domain.** The $\alpha_3$ domain of H-2K$^b$ was folded *in vitro* for the experiments shown in Figure 4C-G. Aliquots were analyzed by nanoscale differential fluorimetry (nanoDSF): the sample is heated in a capillary from 20 °C to 90 °C, and tryptophan fluorescence is recorded every 0.015 K. Unfolding of the protein breaks up the hydrophobic interior of the protein and removes the tryptophan residues (there are four in the $\alpha_3$ domain of H-2K$^b$) from their native interactions, resulting in a change of the local dielectric constant and thus a change of the wavelengths of the fluorescence maxima and of the intensities of the tryptophans, which are read together by the fluorimeter as a summary signal. The first derivative shows as a peak or valley the midpoint of transition, usually called the melting temperature ($T_m$). In the green curves (representing two technical repeats), a clear melting transition of the $\alpha_3$ domain is seen with a $T_m$ of about 54 °C. For the red curves, protein aliquots were heated to 90 °C for 15 minutes before analysis, thus demonstrating the background of unfolded protein in the assay.

#### Supplementary Figure S5

**A**

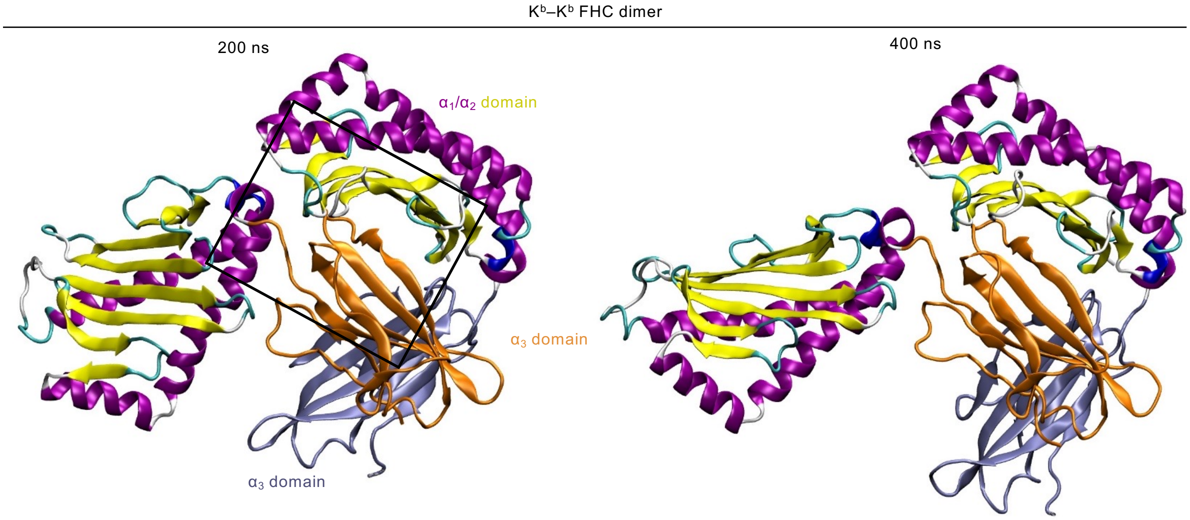


**B**

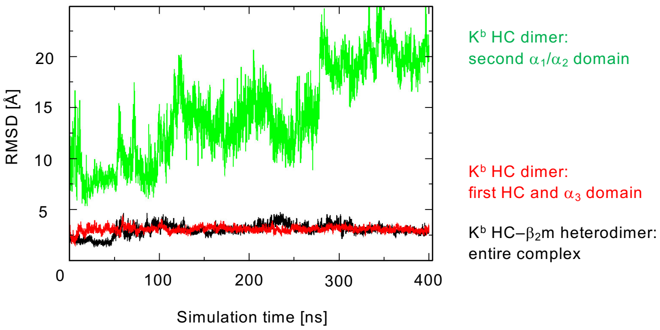


**MD simulation snapshots of an FHC dimer.**

**A**, The $\alpha_3$ domain of one $K^b$ molecule is depicted in orange, the $\alpha_3$ domain of the other $K^b$ molecule is depicted in lavender. The $\alpha_1/\alpha_2$ superdomains are shown in yellow/purple for both FHCs. A favorable interaction of the $\alpha_3$ domain of one $K^b$ molecule (orange) with the $\alpha_1/\alpha_2$ superdomain (yellow/purple) of another $K^b$ molecule (with $\alpha_3$ domain in lavender) remains stable after 200 ns (left) and 400 ns (right). The panels represent the conformations sampled at 200 ns and 400 ns time points indicating the exposure and high mobility of the $\alpha_1/\alpha_2$ superdomain of one of the HCs.

**B**, Root-mean-square deviation (RMSD) of the protein backbone with respect to the starting structure vs. simulation time for the simulation of the $K^b$ HC/$\beta_2$m heterodimer (black line) and for the simulation of the predicted FHC dimer (HC/$\alpha_3$ domain, red line) is low. In the FHC dimer, the relative mobility of the $\alpha_1/\alpha_2$ superdomain of the second HC (green line) after superposition of the trajectory on the first HC is high.

#### Supplementary Movie Legends

**Supplementary Movie S1: Dual-color SMCT identifies FHC association in live cell plasma membranes.** GFP-$K^b$ dimer formation was probed by co-locomotion analysis. Individual GFP-$K^b$ molecules labeled with Rho11 (magenta) and Dy647 (green) in STF1 cells were imaged by TIRF microscopy (scale bar, 2.5 µm; frame rate, 10 Hz; playback speed, 1/3 real time). Movies were acquired in the absence of peptide (FHCs, left), in the presence of peptide (HC/$b_2$m/peptide complexes; center) and in the presence of peptide and a crosslinker consisting of a dimeric anti-GFP nanobody (right). Localized molecules in each channel are encircled, co-locomoting molecules are highlighted by white diamonds.

**Supplementary Movie S2: SMCT demonstrates transient association of FHCs.** Individual GFP-$K^b$ molecules labeled with Rho11 (magenta) and Dy647 (green) imaged by TIRF microscopy (scale bar, 450 nm; frame rate, 10 Hz; playback, 1/3 real time). Selected image section showing SMCT of an individual FHC complex (*i.e.*, in the absence of peptide). Localized molecules in each channel are encircled, co-locomoting molecules are highlighted by white diamonds, with the dissociation occurring between 4.03 s and 4.06 s.

**Supplementary Movie S3:** As Supplementary Movie S2.

**Supplementary Movie S4:** As Supplementary Movie S2.

**Supplementary Movie S5:** As Supplementary Movie S2.
